# Machine learning approaches to assess microendemicity and conservation risk in cave-dwelling arachnofauna

**DOI:** 10.1101/2023.12.19.572471

**Authors:** Hugh G Steiner, Shlomi Aharon, Jesús Ballesteros, Guilherme Gainett, Efrat Gavish-Regev, Prashant P Sharma

## Abstract

The biota of cave habitats faces heightened conservation risks, due to geographic isolation and high levels of endemism. Molecular datasets, in tandem with ecological surveys, have the potential to delimit precisely the nature of cave endemism and identify conservation priorities for microendemic species. Here, we sequenced ultraconserved elements of *Tegenaria* within, and at the entrances of, 25 cave sites to test phylogenetic relationships, combined with an unsupervised machine learning approach to delimit species. Our data identified clear species limits, as well as the incidence of previously unidentified, potential cryptic species. We employed the R package canaper and Categorical Analysis of Neo- and Paleo-Endemism (CANAPE) to generate conservation metrics that are informative for future policy, in tandem with conservation assessments for the troglobitic Israeli species of this genus.

## Introduction

Cave-dwelling taxa are at heightened risk of extinction due to the limited ranges imposed by a single cave system or, in extreme cases, a single cave. These taxa, sometimes referred to as short-range endemics or microendemics, face an outsized threat in the face of disturbance to their habitats and climate change. (Harvey et al., 2011; Mammola et al., 2018). With limited individuals to sample, it is a challenge both to delimit endemic cave species, as well as develop management strategies for endangered taxa (Paquin & Hedin, 2004). Broadly, cave ecosystems share core abiotic features, such as reduction or complete absence of light, high relative humidity, and buffered temperature ranges compared to their surrounding terrestrial surface climates (Barr & Holsinger, 1985). The existence and maintenance of biodiversity in cave habitats is predicated on the ability of biota to adapt to such conditions. Because of this, unique phenotypic changes can be observed in cave-dwelling organisms across the animal tree of life. These changes comprise both reductive features (e.g., atrophy of structures not required for subterranean life), as well as constructive adaptations (e.g., compensatory gains in tactile appendages or olfactory capacity (Re et al., 2018; Riddle et al., 2018)). One of the more conspicuous examples of this phenomenon is the partial or complete loss of eyes in cave-dwelling species. The Mexican cavefish (*Astyanax mexicanus*) is a well-studied exemplar of eye loss in cave-dwelling species. The blind morph of *A. mexicanus* is said to have evolved as recently as 20,000 years ago, exemplifying phenotypic change over rapid timescales and without the requirement of reproductive isolation (Fumey et al., 2018). Rapid evolution of disparate phenotypes allows for the study of how speciation begins in cave populations versus surface populations. Over time, speciation may establish troglobites, or species that are obligate cave-dwellers; these can occur in proximity to epigean (surface) counterparts or to troglophiles, species that are facultative cave-dwellers (Howarth, 1983). Because of this, inferring the phylogenetic relationships between closely related troglobites and troglophiles presents a unique challenge.

The Levant, in particular, harbors high diversity of cave fauna in a small geographic area (Gavish-Regev et al., 2021; Peel et al., 2007), due to the incidence of 3 distinct climate zones (Mediterranean climate, semi-arid, desert; *sensu* Köppen-Geiger climate classification system) at the margin of 3 continents. Climatic shifts during the Pleistocene saw the Southern Levant act as an unglaciated refuge for species, with animals colonizing the region from surrounding glaciated areas (Tchernov & Belmaker, 2004). With an abundance of geologically diverse caves, Israel is an ideal locale for the study of phenotypic evolution and speciation of endemic groups in cave ecosystems. Recent surveys of cave sites around Israel have yielded the discovery of several microendemic spider species with troglobite and troglophile representatives (Gavish-Regev et al., 2021). These spiders exhibit a spectrum of eye loss ranging from “normal,” fully developed eyes to complete absence of eyes.

In a recent work, 7 new species of troglobitic *Tegenaria* were described using traditional morphological approaches, with barcoding (COI sequencing) and ddRAD sequencing reinforcing interpretations of species boundaries (Aharon et al., 2023). The Israeli troglobitic species were recovered as a distinct clade from local troglophilic species. These data suggested that the current cave species are relicts that descended from a single, surface-dwelling ancestor. While these data supported species recognition and a hypothesis for the evolutionary history of this group, interspecies relationships were disputed across the 2 different sequencing methods, possibly due to high locus drop-out rates in RAD sequencing across previously unrecognized species boundaries. The resulting data matrices thus bore few sites that were present across all taxa, a known driver of phylogenetic uncertainty (Roure et al. 2013).

Here, based on a target-capture sequencing approach that leverages ultraconserved elements,, we inferred a phylogeny to resolve these relationships, applied an unsupervised machine learning approach to delimit species, and assessed conservation priority of Israeli *Tegenaria.* Considering the ensuing inferences of species boundaries, we generated metrics of phylogenetic diversity to identify areas of unanticipatedly high endemism across Israeli cave sites that warrant conservation priority. We then classified the conservation status of each troglobitic species according to the IUCN Red List Categories and Criteria workflow.

## Materials and Methods

### Species sampling and sequencing

We sequenced samples from 161 *Tegenaria* specimens freshly collected from field campaigns across 25 caves of Israel (2018-2020). Outgroup taxa consisted of representatives of the confamilial genera *Agelena* (N = 4) and *Lycosoides* (N = 4). Samples were extracted from specimens using a Qiagen DNeasy Blood & Tissue Kit and eluted in 10 mM Tris-HCl, with additional bead-based purification of previously generated EDTA-preserved extractions (from Aharon et al. 2023). DNA was quantified using a Qubit 3 fluorometer with a High Sensitivity dsDNA Assay Kit. Enzymatic fragmentation, end repair/A-tailing, adapter ligation, and library amplification were performed with a KAPA HyperPlus Kit and dual index primers (i5 and i7) for multiplex sequencing. Additional details of the library preparation procedures are provided in S3.

Samples were pooled samples by groups of 8, each with 125 ng of DNA for subsequent target capture, following the myBaits protocol v5.02 (Arbor Biosciences) for targeted enrichment using a spider-specific probe set (Kulkarni et al., 2020). Pools were sequenced at the University of Wisconsin Biotechnology Center on an Illumina NovaSeq 6000 2 · 150 bp S1 flow cell. Cleaning and trimming of raw read data were performed using illumiprocessor v2.0, followed by assembly of libraries using AbySS 2.0. We employed the software package PHYLUCE v1.6 for subsequent data processing and analysis (Faircloth, 2015). Individual scripts used through PHYLUCE are indicated in S3.

### Phylogenomic analysis

To assess sensitivity to data completeness, we applied successive gene occupancy thresholds of 50% (777 loci, 161 taxa), 90% (227 loci; 161 taxa), and 95% (88 loci; 161 taxa). Concatenation-based maximum likelihood analyses were performed using IQTREE v.2 (Nguyen et al., 2014). Automated model selection was performed using ModelFinderPlus (Hoang et al., 2018; Kalyaanamoorthy et al., 2017). Nodal support was estimated via 1500 ultrafast bootstrap replicates and 1500 bootstrap replicates for the SH-like approximate likelihood ratio test (Guindon et al., 2010; Hoang et al., 2018). For phylogenetic analyses using multispecies coalescent methods, species trees were estimated with ASTRAL v. 3, using individual gene trees as inputs, with gene tree topologies estimated using IQ-TREE v. 2 under automated model selection.

To test the robustness of the ensuing tree topologies, we additionally assessed phylogenetic signal across UCE loci using *genesortR*. This method, which implements a principal components-based approach to quantifying phylogenetic informativeness, requires a resolved species tree *a priori* for the computation of Robinson-Foulds distances for each gene tree. To limit the influence of unresolved parts of the species tree on the ranking of phylogenetically useful genes, we collapsed 2 nodes in the species tree that were not resolved with maximal nodal support. Subsequent to sorting on phylogenetic signal, we inferred the maximum likelihood tree topology from a concatenated matrix comprised of the 100 highest-ranked and informative loci.

### Machine learning-based validation of species boundaries

To independently validate the species limits suggested by our phylogeny, we employed a variational autoencoder (VAE), an unsupervised machine learning technique, which was previously demonstrated to be informative for species delimitation in a cryptic genus of harvestmen (Derkarabetian et al., 2019). To implement VAE, we first called single nucleotide polymorphisms (SNPs) from our UCE data using the Genome Analysis Toolkit (McKenna et al., 2010). These data were then encoded in the one-hot format, which allows for the representation of categorical data as vectors with integer values. In the case of sequence data, the vector [1, 0, 0, 0] represented an A, [0, 1, 0, 0] was C, [0, 0, 1, 0] was G, and [0, 0, 0, 1] was T. Importantly, sites where data was missing were encoded as [0, 0, 0, 0] which prevented the model downstream from erroneously grouping terminals solely based on shared missing data. Following the encoding step, VAE was implemented using the TensorFlow (Abadi et al., 2016) and Keras python libraries. A script by Derkarabetian et al. (2019) was used to build the model and plot the results.

### CANAPE

To assess the conservation priority of Israeli *Tegenaria* we conducted Categorical Analysis of Neo- and Paleo-Endemism (CANAPE) on our full dataset (Mishler et al., 2014). We used canaper, a package which allows for the running of CANAPE in its entirety in R (Nitta et al., 2023). As inputs for canaper, we used our 50% taxon occupancy phylogeny and a data frame consisting of a column with each terminal name and 2 columns with the latitude and longitude from which the corresponding samples were collected. To be read by canaper, the locality data frame had to be converted to a community matrix which was further converted to a tibble. The first step of the CANAPE analysis was to run *cpr_rand_test* which generates a set of random communities with a series of metrics to which input data is compared. For our purposes, we opted for 500 random communities and used the null model “curveball” for randomization (Strona et al., 2014). The next step was to run the *cpr_classify_endem* function to classify each of our communities as paleoendemic, neoendemic, or both. We then visualized these data as a map of our study site with color coded grid cells corresponding to the endemism type.

### Conservation assessments and IUCN designation

To perform conservation assessments of troglobitic Israeli Tegenaria, we used data from the Israel National Arachnid Collection, where Tegenaria specimens from our cave surveys were deposited, and previous studies we have published on this system (Aharon 2023, Gavish-Regev 2021). Distribution records of all Tegenaria species were extracted from original descriptions (Aharon et al 2023), as well as from our ecological surveys. To generate measurements of Extent of Occurrence (EOO) and Area of Occupancy (AOO) we performed spatial analysis using the R package red (Cardoso 2017).

## Results

### Relationships of Tegenaria inferred from UCE-based datasets

Maximum likelihood analyses recovered 2 robustly supported clades of Israeli *Tegenaria*, corresponding to a clade of eye-bearing epigean species and a clade of species with eye reduction or complete loss (Fig. 2). Relationships between species were robustly supported across analyses (bootstrap resampling frequency [BR] >99%), barring the placement of *Tegenaria trogalil* (BR = 62%). Whereas the epigean clade exhibited deep genetic distances between species (i.e., long patristic distances between clades), the base of the troglobitic clade exhibited short branch lengths, potentially corresponding to a rapid radiation. Within the troglobitic clade, species inhabiting caves in the north of Israel formed a grade with respect to a nested clade of southern species (BR = 99.9%).

**Figure 1.**
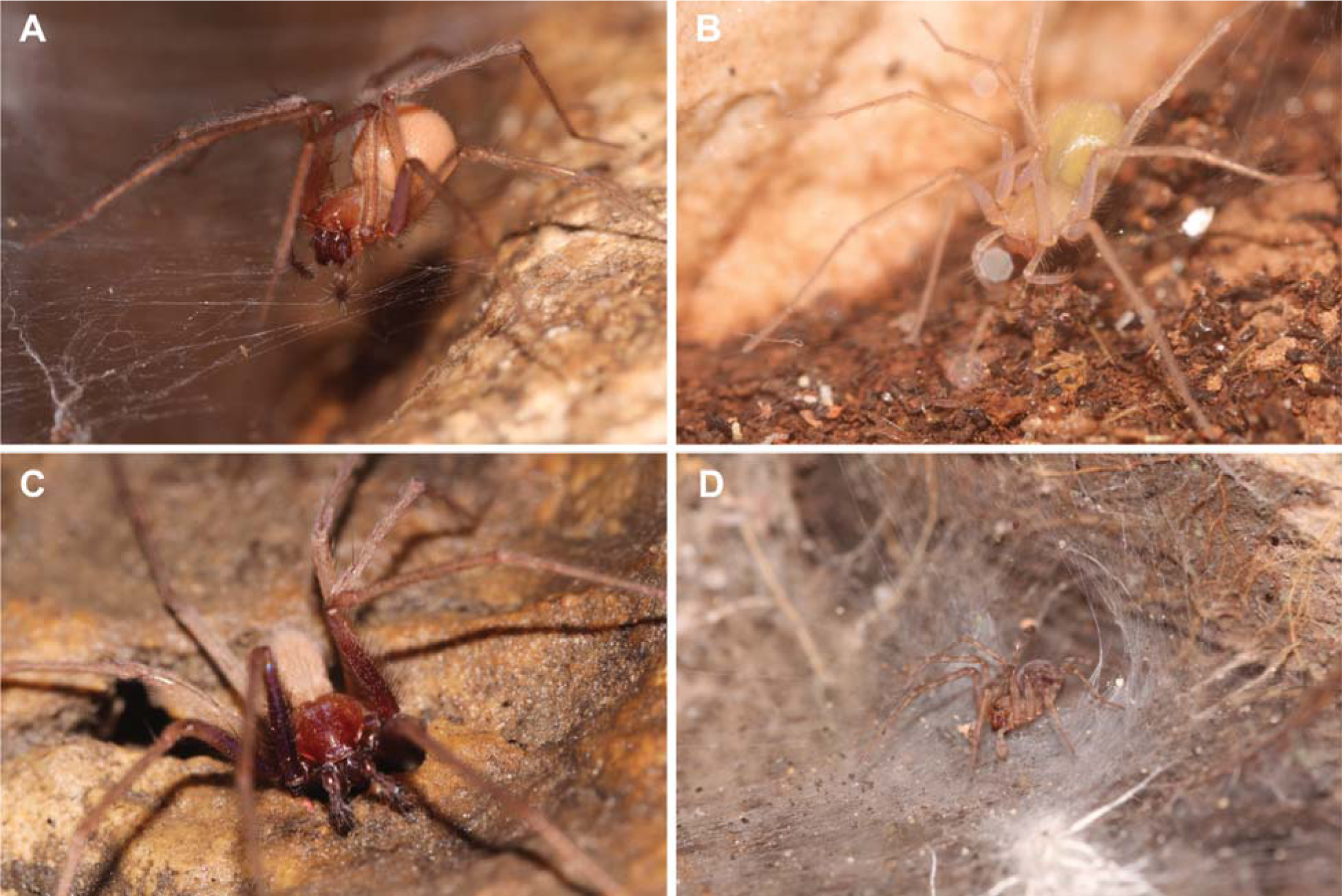
Live habitus of Israeli Tegenaria. **A.** Eyeless T. ornit from dark zone of Ornit Cave. **B.** Eyeless T. naasane from ‘Arak Na’asane Cave. **C.** T. yaaranford from with reduced eyes from the dark zone of Te’omim Cave. **D.** The epigean T. pagana with fully developed eyes at the entrance of Te’omim Cave. All pictures by S. Aharon.

**Figure 2.**
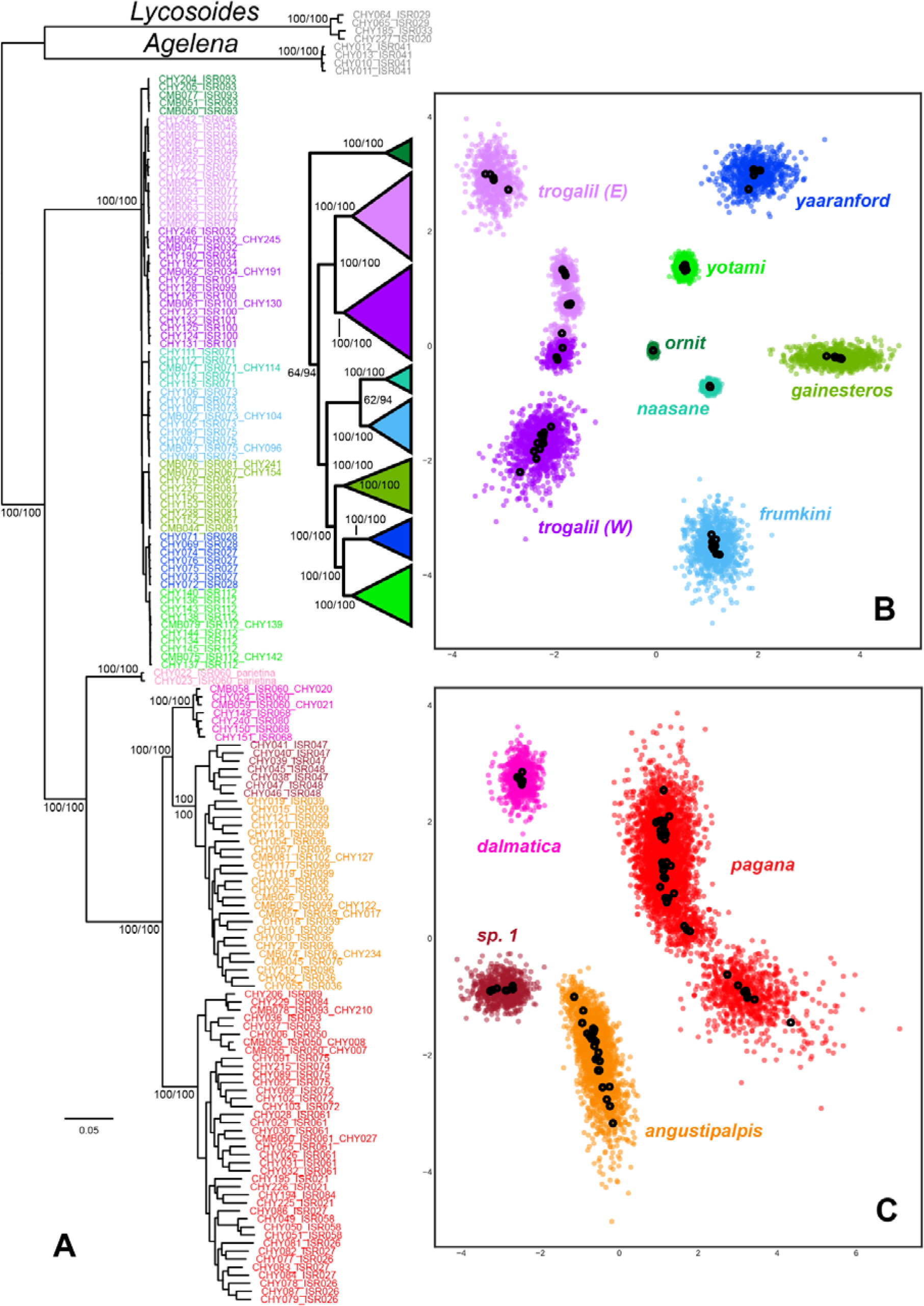
Phylogeny and inferred species groups of Israeli *Tegenaria* based on UCEs and unsupervised machine learning approach (VAE). **A.** Maximum likelihood phylogeny of 161 terminals based on 777 UCE loci. The upper group (cool colors) corresponds to the troglobitic clade, and the bottom group (warm colors) corresponds to the epigean clade. **B.** Visualization of VAE analysis of troglobitic clade. Black circles represent the mean position of individuals and colored circles represent standard deviations. Note the complexity of the *T. trogalil* species cluster, whose standard deviations overlap as a third, separate cluster. **C.** Visualization of VAE analysis of epigean clade. Note the detection of an undescribed species.

### VAE-based inference of species clusters

We used VAE to delimit species within Israeli *Tegenaria*, with an emphasis on delimiting the seven recently described troglobitic species. For computational efficiency, species delimitation was performed separately for each clade, with one corresponding to the epigean clade and one corresponding to the troglobitic clade. Broadly, VAE inferred population clusters corresponding to species inferred by the phylogeny (Fig. 3). In the epigean clade, clusters were well-defined and exhibited no overlap in standard deviation. The *T. pagana* cluster appeared discontinuous but distinct from the rest of the clusters. In the troglobitic clade, species limits were again well-defined, with the notable exception of *T. trogalil* species, which resolved as three separate clusters. 2 of these corresponded to the East and West Galilee caves and a third corresponded to a cluster comprised of both regions. There was also a tendency for VAE to group some of the reduced eye species from more isolated caves (e.g., *T. ornit, T. naasane*) into tighter clusters (i.e., low variance).

**Figure 3.**
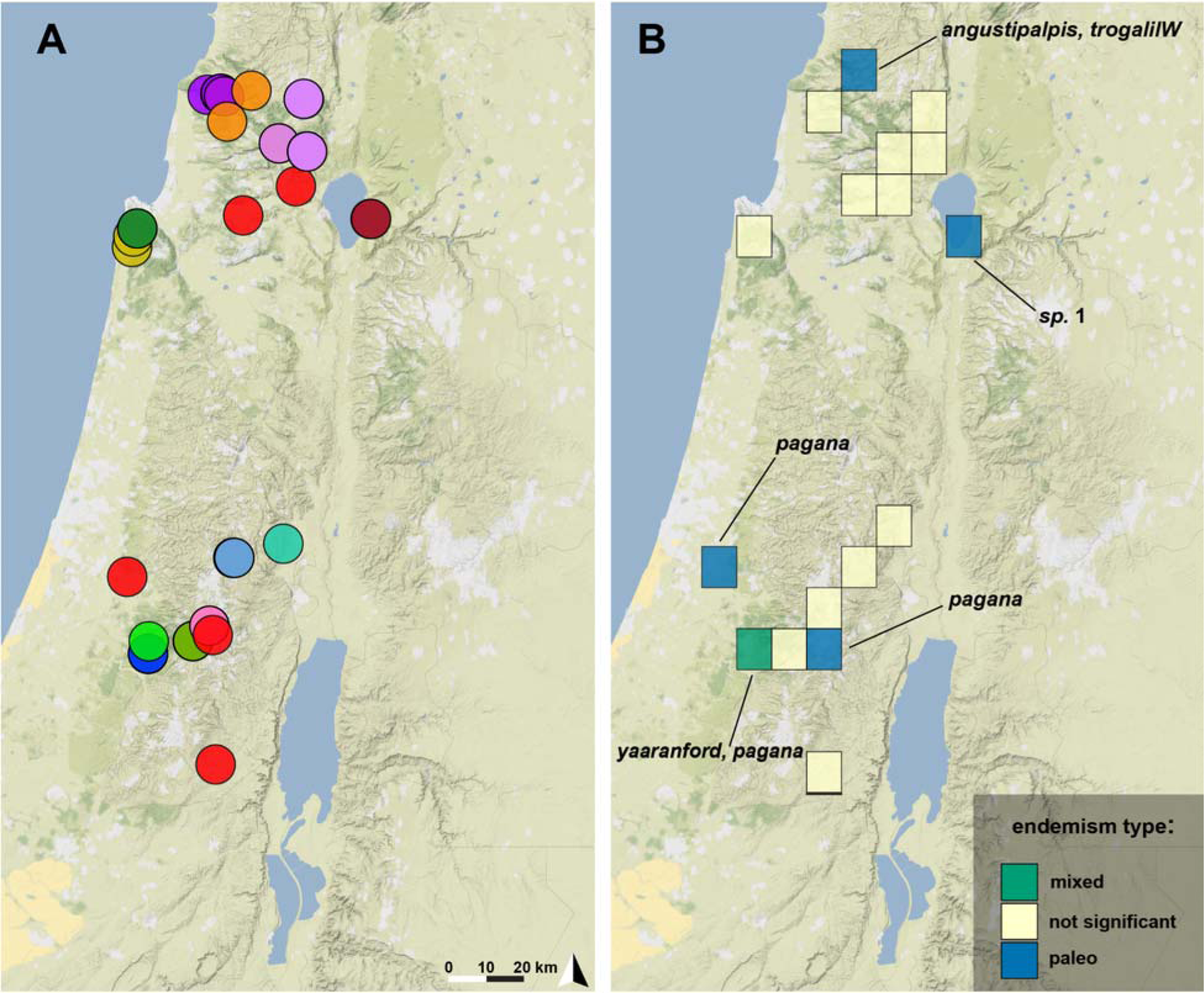
Maps of localities and CANAPE analysis. **A.** Map of localities from which all specimens used for sequencing were collected. Colors correspond to colors of tip labels in phylogenetic tree and VAE figures. **B.** Results from CANAPE analysis. Boxes are 10 km^2^ grid cells that encompass all localities sampled. Note the four sites of paleo-endemism and single site of mixed endemism. The sites of paleo-endemism are home to epigean species with the exception of T. trogalil (West). The site of mixed endemism is home to Te’omim cave which harbors the troglobitic T. yaaranford and epigean T. pagana species.

### Identification of biodiversity hotspots across Levantine cave sites

5 caves out of the 25 surveyed harbored a single species found in no other caves. Out of 17 10km^2^ grid cells in our study area, 4 were classified as sites that exhibit paleo-endemism and one was classified as mixed (paleo- and neo-endemism). The species represented in the paleo-endemic sites are *T. pagana, T. trogalil* (West caves)*, T. angustipalpis*, *T. sp. 1* (Fig. 4). One such species, *T. trogalil* from Namer cave, is the sole representative of the troglobitic clade that occurs in a site of paleo-endemism.The single site of mixed endemism is home to *T. pagana* (fully developed eyes) and *T. yaaranford* (reduced eyes) at Te’omim Cave (Fig, 4).

### Conservation assessment using IUCN workflow

Upon evaluating the conservation status of the 7 troglobitic *Tegenaria* species found in Israel, we classified five as Critically Endangered (CR) due to extremely small EOO and AOO, as well as isolation and very low population size (<50 mature individuals) observed in some of the caves. One troglobitic species that is found in several (likely connected) neighboring caves was classified as Endangered (EN), due to threats of development that have the potential to destroy its habitat. We classified only one species as Vulnerable (VU) due to projections of habitat extent and quality decline, as we observed a low number of mature individuals distributed across six sites.

## Discussion

As global biodiversity continues to diminish due to rapidly changing climatic conditions and human factors, the conservation of species has never been more important. Currently, 41,000 species are threatened with extinction (International Union for Conservation of Nature, 2022). If biodiversity loss persists at an increasing rate, the stability of ecosystems is in jeopardy. Notably, conservation status statistics published by the International Union for Conservation of Nature (IUCN) only reflect species that have been assessed and therefore may not serve as an accurate representation of the conservation status of many small-bodied and enigmatic taxa. For instance, the order Arachnida is proportionately the least represented animal taxon that is currently assessed by the IUCN, with only 0.4% of described species assessed (IUCN Red List version 2022-2: Table 1a). Therefore, it is imperative that specific efforts be made to assess the conservation status and priority of underrepresented groups such as arachnids.

Assessing the conservation status of spiders is difficult due to insufficient taxonomic and biological information on many lineages as well as difficulties in evaluating population size. Few long-term monitoring schemes are in place and often data on species occurrences is anecdotal or from a single sampling event in a specific locality. Since we have been studying and monitoring spiders in caves in Israel for over a decade, these efforts have enabled us to gain knowledge on the distribution of spiders in caves in Israel. The result that five out of seven new species described are classified as CR is unsurprising, given their small population sizes and limited range.

Short-range endemic (i.e., microendemic) species are highly vulnerable to environmental degradation and anthropogenic disturbance. Effective recognition and conservation assessment of short-range endemics requires a combination of taxonomy and genomic datasets for phylogenetically-informed assessment of biodiversity metrics (Agnarsson & Kuntner, 2007; Harvey et al., 2011; Mishler et al., 2014). Here, we deployed unsupervised machine learning for identifying species limits, prior to assessing conservation priority for several microendemic spiders of Levantine cave habitats. Our results suggest that troglobitic Israeli *Tegenaria* are both genetically isolated from each other and distantly related to their surface-dwelling relative counterparts. Contrary to established models of cave adaptation (i.e., repeated colonization of caves by epigean taxa), the Israeli *Tegenaria* comprise a unique evolutionary case wherein troglobitic species likely resulted from a rapid radiation of a single ancestral species that had a proclivity for cave-blindness, with possible extirpation of their epigean ancestor. The epigean species found in Israel today are more closely related to Mediterranean *Tegenaria*, suggesting a second wave of colonization of shallower parts of caves in Israel (Aharon et al., 2023).

The clustering proposed by VAE corroborated species limits inferred by our phylogenetic analyses. However, it also detected clusters not recovered as species in our phylogeny, suggesting that there may be yet undescribed species, both within the cave-dwelling and the surface-dwelling clades. In the case of the troglobitic clade, we observed a new cluster within *T. trogalil* comprised of exemplars of both the eastern and western caves, whereas standard phylogenetic analyses tended to recover the eastern and western caves as separate and monophyletic groups. The detection of this mixed cluster within *T. trogalil* could represent complex dynamics stemming from introgression or incomplete lineage sorting. Within the eye-bearing clade, VAE detected a possible new cluster within *T. pagana*, which corresponds to specimens from five localities from the north of Israel. This lineage was recovered as the sister group to the remaining *T. pagana* and may constitute a new cryptic species.

Taken together, the high regional fidelity and deep genetic distances between species suggest that Israeli caves serve as both “museums” of ancient diversity and “cradles” of recent diversification for Levantine arachnofauna. Single cave sites can be host to faunal antiquity from 2 separate sources (epigean and troglobitic), resulting in high metrics for paleoendemism. However, biodiversity metrics like those implemented for CANAPE are grounded in the expectation that species in the occurrence matrix are neither too rare, nor too widespread. The value of microendemic species is difficult to capture using algorithmic approaches to biodiversity assessment, because sites with a single microendemic species that harbors low genetic diversity (e.g., *T. ornit*) are not highly ranked for metrics like phylogenetic diversity or phylogenetic endemism. Moreover, sites that harbor a microendemic of one taxon may harbor other such endemics that are outside of the focal study system. As an example, Ornit cave, which has one of the smallest dark chambers of all the caves surveyed, harbors the entirely blind species *T. ornit*, as well as an undescribed species of a palpigrade (microwhip scorpion) and a pseudoscorpion. The use of software like CANAPE alone to assess conservation priorities thus comes with a tradeoff.

Our results substantiate the need for regulations surrounding access to, and conduct within, caves in Israel by tourists and visitors. Moreover, we suggest that abiotic conditions as well as populations in those caves will be monitored, to assess any changes in population size. It is imperative that the public is made aware of the extreme rarity and uniqueness of the species within the Israeli *Tegenaria* system. We recommend that caves containing type-locality endemics be identified with signage indicating to visitors that if they intend to enter the cave, they must use extreme care when maneuvering about the cave to not disturb any spiders, webs, or egg sacs they may encounter. Only some of those type-locality caves are protected by Israeli law, as they are located in nature reserves (e.g., Ornit, Teomim, Sarah and Namer caves) or national parks (Soreq cave), while others are not protected (e.g., En Sarig spring tunnel, Yir’on cave). But even the caves that are protected host many visitors that affect abiotic conditions, or even actively alter microhabitats by lighting fires inside the caves. Broadly, all these caves should be identified by appropriate government agencies as sites of natural historical significance and receive protections commensurate with this status, that will forbid use of fire inside the cave and close the deep chambers for visitors. With 5 out of the 7 new species classified as CR under IUCN guidelines, it is imperative that these regulations are put in place with deliberate haste.

