## Supplementary Figures for "Machine learning approaches to assess microendemicity and conservation risk in cave-dwelling arachnofauna"

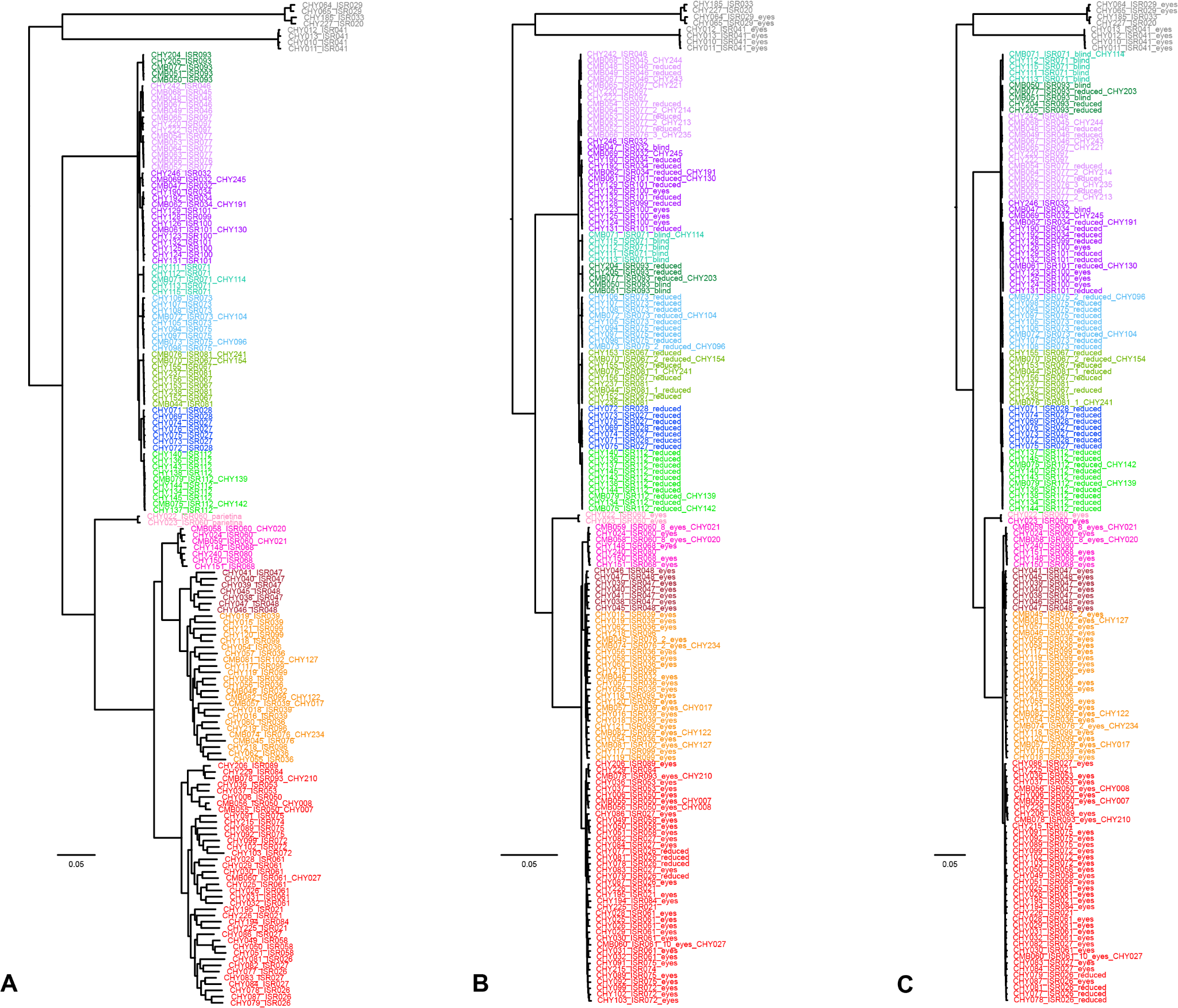

**Figure S1.** Maximum likelihood analyses at different gene occupancy thresholds using IQTREE v.2. All analyses sampled the same 161 terminals. **A.** 50% gene occupancy tree based on 777 UCE loci. This tree was used as input for VAE and CANAPE analyses. **B.** 90% gene occupancy tree based on 227 UCE loci. Note the change in placement of *T.* *ornit* (dark green) which was recovered as nested within the troglobitic clade. **C.** 95% gene occupancy tree based on 88 UCE loci. Note the changes in placement of *T.* *ornit* (dark green) and T. *Naasane* (turquoise) which were recovered in a clade sister to the rest of the troglobitic species.

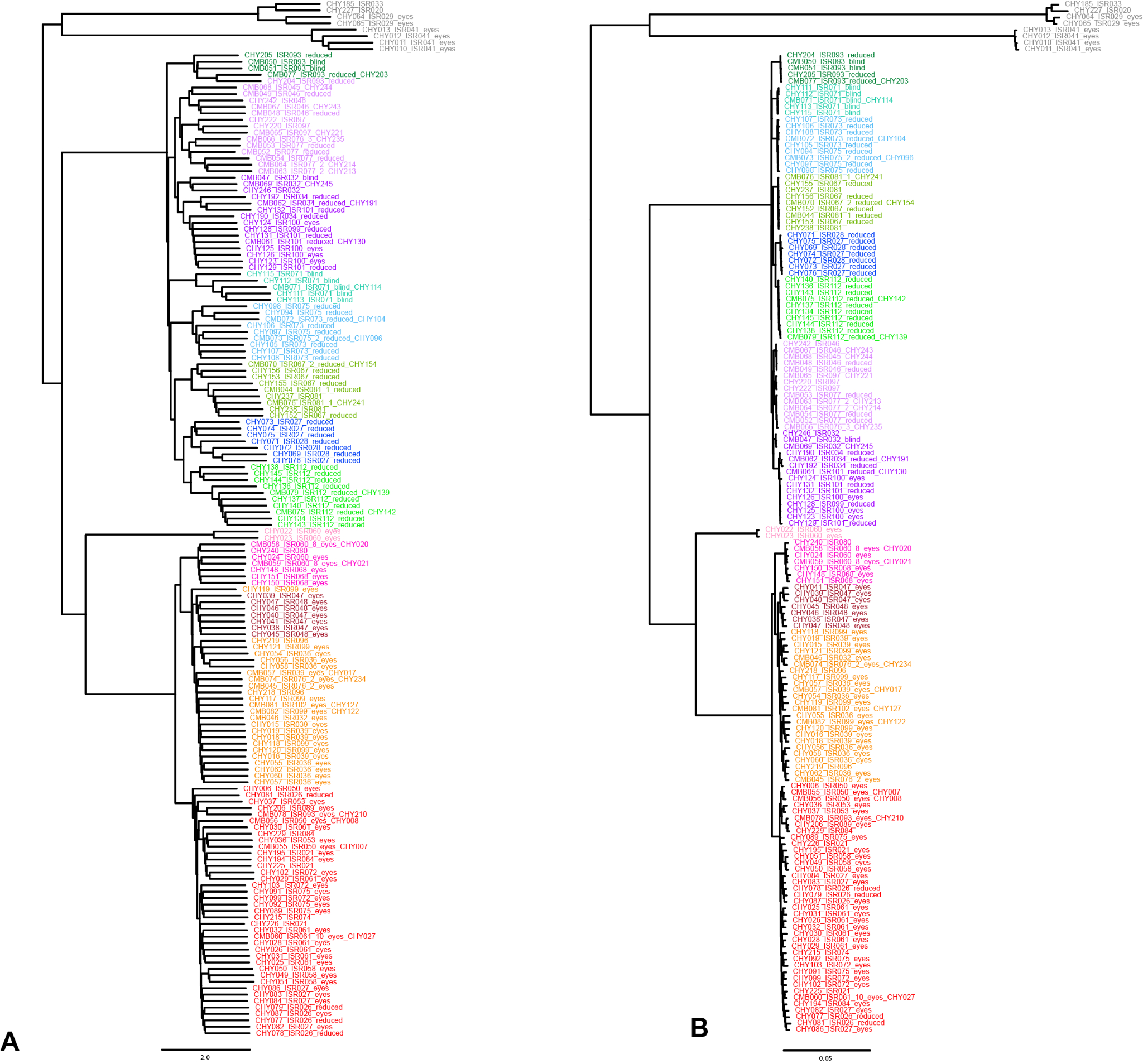

**Figure S2.** Gene tree and phylogenetic robustness analyses. **A.** Species tree implemented using ASTRAL v.3 based on individual gene trees from IQTREE analyses of 777 loci data matrix. Tree topology was conserved with the exception of the yet undescribed *Tegenaria* sp. 1, which was recovered within *T*. *angustipalpis*. **B.** Tree based on 100 most phylogenetically informative loci from *genesortR* analyses. Note the placement of the *T. trogalil* clade more deeply within the rest of the troglobitic clade compared to maximum likelihood analyses.
