## Supplemental File 1 for "Machine learning approaches to assess microendemicity and conservation risk in cave-dwelling arachnofauna"

**Assessment workflow:**

Here, we evaluated the 7 troglobite *Tegenaria* species found only in caves in Israel and Palestine. We classified five troglobite species, that are each found in a single cave locality, as Critically Endangered (CR) due to very small EOO and AOO, isolation, and low population densities. One troglobite species found in several neighboring caves, likely connected in the epikarst, was identified as Endangered (EN) due to threats of development that could destroy their habitat. We identified only one species as vulnerable (VU) due to projection of habitat extent and quality decline as well as a low number of adults observed. Moreover, there is a possibility that this species found in 6 caves in the Galilee is a complex of cryptic species.

1. *Tegenaria frumkini* Aharon & Gavish-Regev, 2023

**Assessment Rationale**

This is an endemic cave-dwelling funnel-web spider, found only in the twilight and deep zones of a few caves in the Ofra karst basin, located in Samaria mountains, Israel (West Bank)(Frumkin & Langford, 2017). Extensive surveys in other caves outside of the Ofra karst basin have been conducted and the species was not found elsewhere. The abiotic conditions in the cave are fragile and may be threatened by climate-related changes and extreme weather events such as drought, heat, in addition to extensive urban development in the karst basin. The habitat extent and quality, and the number of adults is therefore projected to decline in the near future, making it Endangered under Categories B1 and B2.

**Species information**

*Tegenaria frumkini* Aharon & Gavish-Regev, 2023

**Common names**

Cave funnel-spider

**Taxonomy**

**Kingdom	Phylum	Class	Order	Family**

Animalia	Arthropoda	Arachnida	Araneae	Agelenidae

**Taxonomic notes**

This troglobitic species is found in the twilight and deep zones of a few neighboring caves in the Ofra karst basin, in the Samaria mountains, Israel (West Bank) (Aharon et al., 2023, p. 20). Males, females and juveniles of this spider species are depigmented and bare eight reduced eyes, (Figure 1).

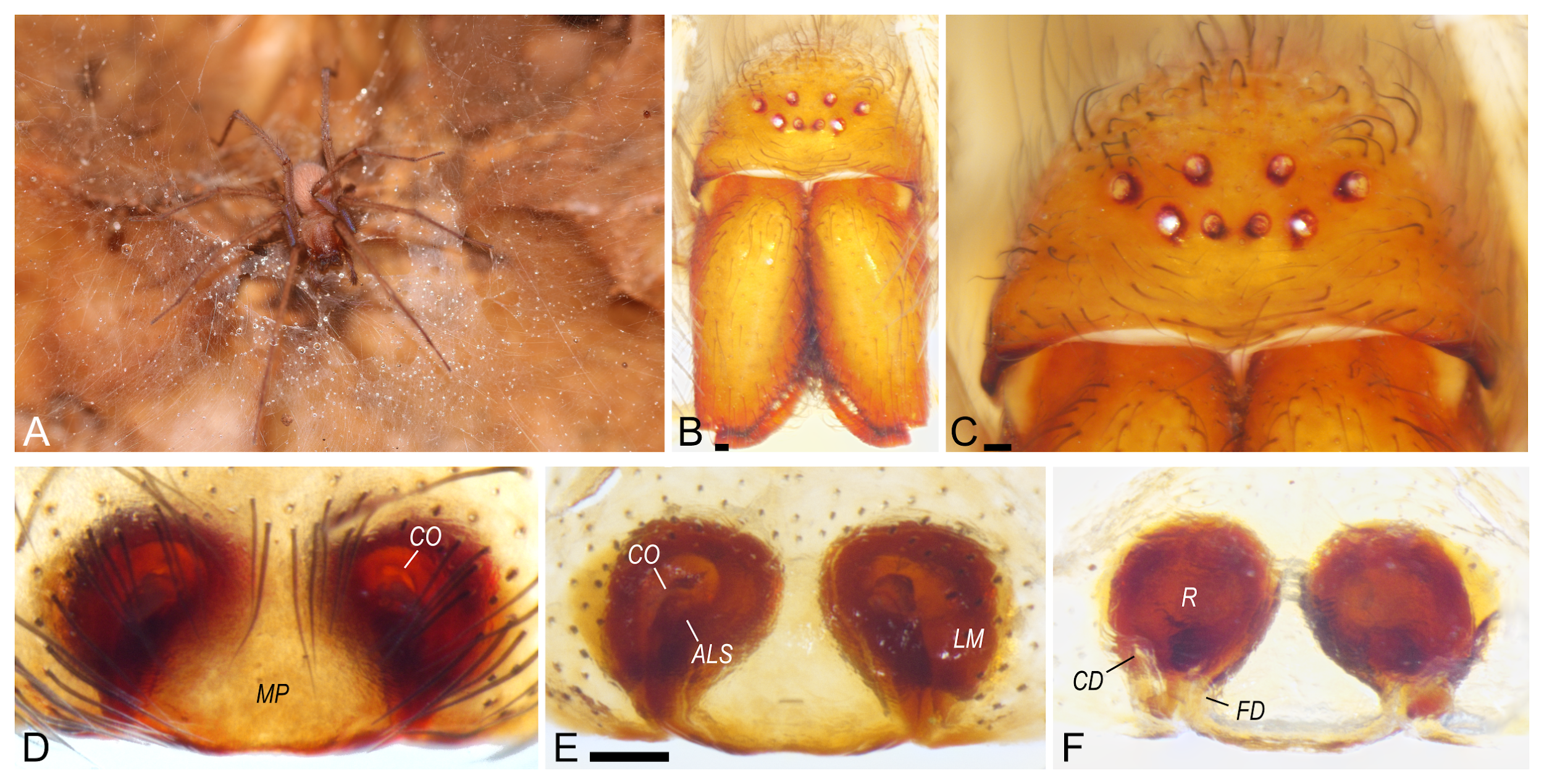

**Figure 1:** *Tegenaria frumkini* Female, Ofra caves. Shlomi Aharon.

**Region for assessment:** Global

**Biogeographic realm:**  Palearctic, Levant

**Countries:** Israel (West Bank)

**Map of records:**

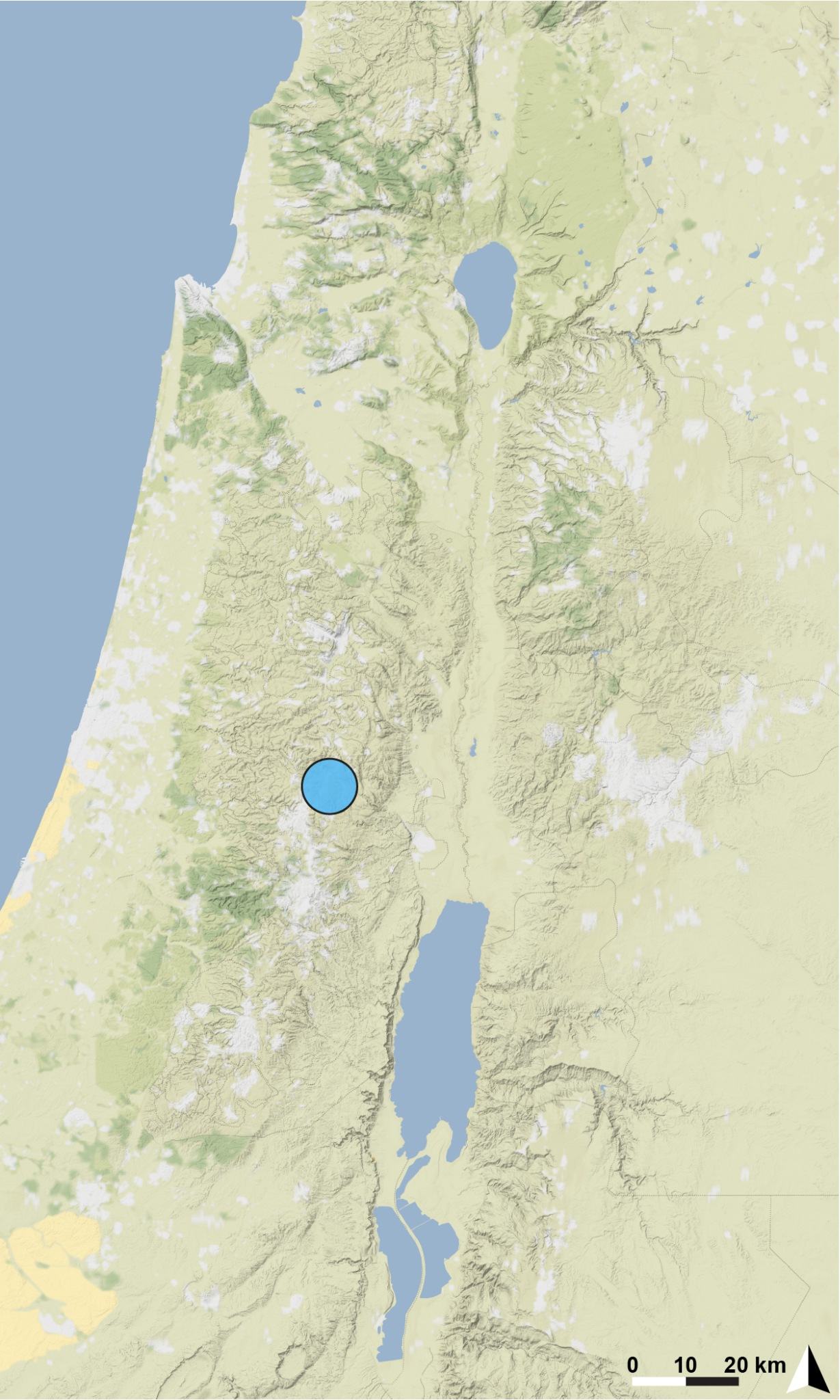

**Basis of EOO and AOO:** Observed

**Basis (narrative)**

Although more than 100 caves in Israel and Palestine were surveyed intensely for arachnids since 2012, the Ofra karst field caves are the only locality known for this troglobitic *Tegenaria* species.

**Range description**

This is an endemic species restricted to the twilight and deep zones of a few neighboring caves in Ofra karst basin, Samaria.

**Extent of occurrence (EOO) estimate(km^2^):** <1km^2^

**Trend:** Unknown.

**AOO (km^2^):** 8

**Trend:** Unknown.

**Number of locations:** 6

**Justification for number of locations:** only found in six geographically neighboring caves despite extensive surveys through the region.

**Habitat**

**System:** terrestrial

**Habitat specialist:** Yes

The species is a specialized troglobite living in the twilight and deep zones of a few geographically neighboring caves in Ofra karst basin (5 km^2^), where temperature and humidity conditions are constant.

**Trend in extent, area or quality?:** Yes

**Justification for trend:**

Humidity and temperature change, changes in abiotic conditions due to extreme events such as global warming. This species is found only in the karst basin of Ofra, and urban development will potentially destroy the caves, which could wipe out the species.

**Habitat importance:** Major Importance

**Habitats:**

- 7. Caves and Subterranean Habitats (non-aquatic)

- 7.1. Caves and Subterranean Habitats (non-aquatic) – Caves

**Ecology**

**Size:** 6-10 mm

**Generation length (yr):** unknown

**Ecology and traits (narrative)**

The species builds funnel-webs in small holes in rock walls of caves, as well as under and near stones in the cave. Found only in the twilight and deep zones (Aharon et al. 2023).

**Threats:**

These caves are now under threat due to urban development in the Ofra karst basin. This species is found only in the Ofra karst basin, and one urban development project could destroy these caves, which could wipe out the species. In addition, climate change is predicted to affect the entire karst basin abiotic conditions, extreme weather events such as drought and high temperatures could all destroy the cave conditions in just one event and will have a severe effect on this micro-predator.

**Ongoing**

-  1. Residential & Commercial Development

-  1.1 Housing & Urban Areas  – Urban development

- 6. Human intrusions & disturbance

- 6.1. Human intrusions & disturbance – Urban development

- 11. Climate Change & Severe Weather

- 11.2 Extreme droughts

- 11.3 Temperature extremes

**Conservation**

The cave is not protected by law, and there are plans for urban development in the karst field, where the caves are located. The caves should be protected from urban development.

**Research needed:**

- 3. Monitoring

- 3.1. Monitoring - Population trends

- 3.4. Monitoring - Habitat trends

Endangered (EN): B1a,b(iii)+B2a,b(iii)

2. *Tegenaria gainesteros* Aharon Gavish-Regev, 2023

**Assessment Rationale**

This is an endemic funnel-web spider, found only in a single spring water tunnel, in the Judean mountains. Extensive surveys in other water tunnels and caves have been conducted and the species was not found elsewhere. The abiotic conditions in the water tunnel are fragile and may be threatened by climate-related changes and extreme weather events such as drought and heat. In addition conditions are threatened by anthropogenic effects such as fire in the vicinity of the spring. Furthermore, this spring is very touristic during summertime. The habitat extent and quality, and the number of adults are therefore projected to decline in the near future, making it Critically Endangered under Categories B1 and B2.

**Species information**

*Tegenaria gainesteros* Aharon Gavish-Regev, 2023

**Common names**

Cave funnel-spider

**Taxonomy**

**Kingdom	Phylum	Class	Order	Family**

Animalia	Arthropoda	Arachnida	Araneae	Agelenidae

**Taxonomic notes**

This troglobitic species is found in a single spring water tunnel, mainly in the dark parts, in ’En Sarig, the Judean mountains in Israel (Aharon et al., 2023) . Males, females and juveniles of this spider species are depigmented and bare eight reduced eyes, (Figure 2).

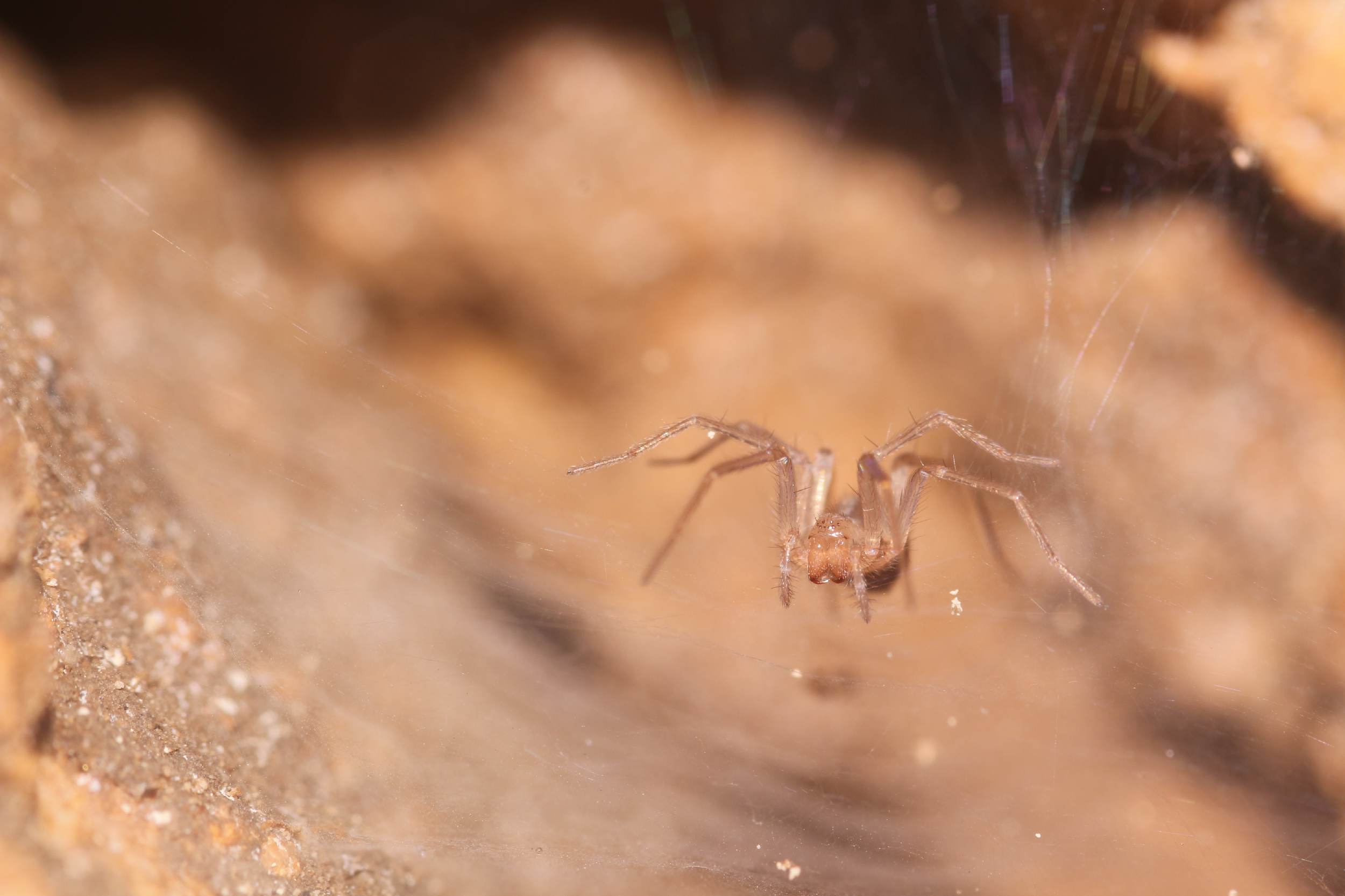

**Figure 2:** *Tegenaria gainesteros* Female, En Sarig water tunnel. Shlomi Aharon.

**Region for assessment:** Global

**Biogeographic realm:**  Palearctic, Levant

**Countries:** Israel

**Map of records:**

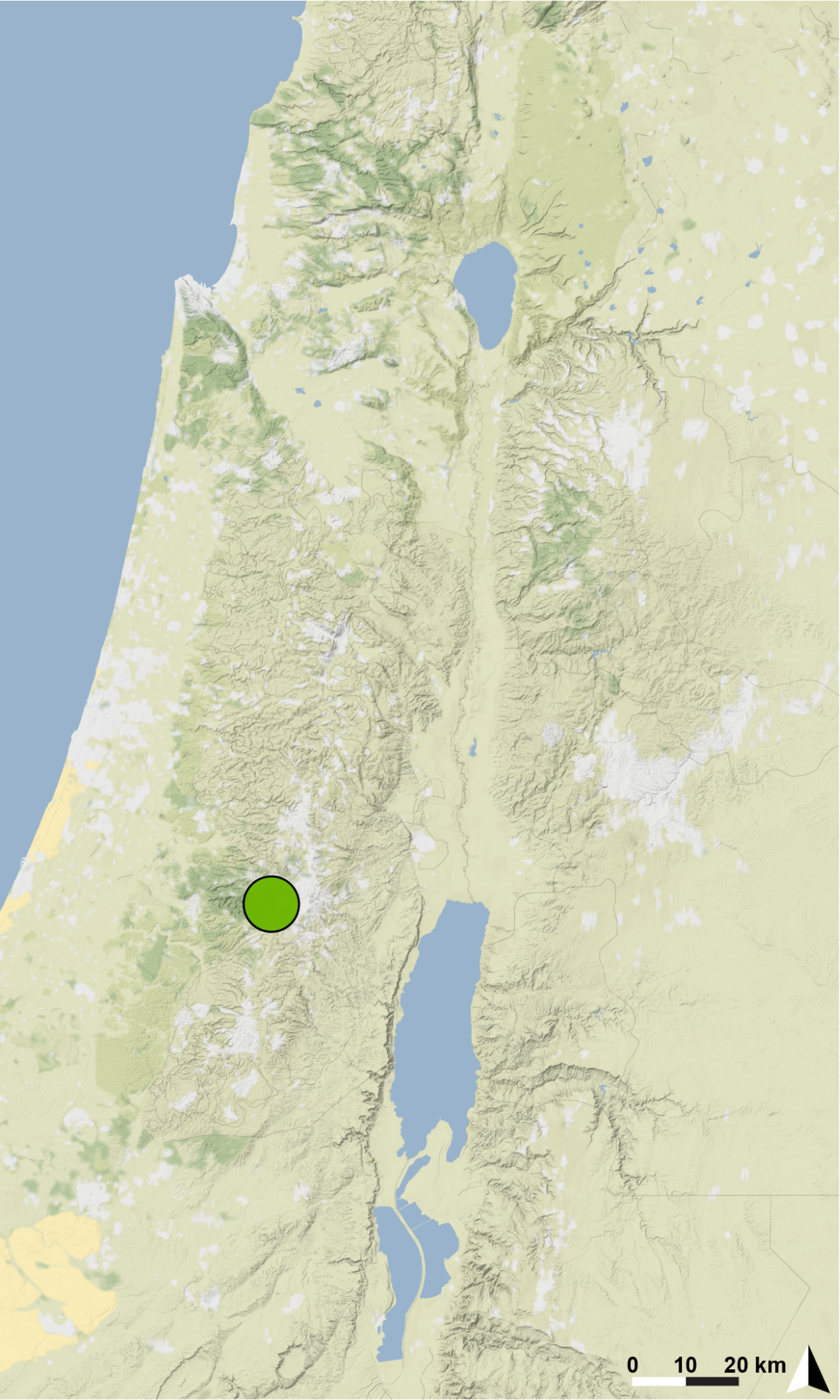

**Basis of EOO and AOO:** Observed

**Basis (narrative)**

Although more than 100 caves in Israel, including several water tunnels the Judean mountains, were surveyed intensely for arachnids since 2012, ’En Sarig is the only locality known for this troglobitic *Tegenaria* species. It is possible that this species will be found in a similar habitat if monitoring is conducted in the spring tunnels of the Judean mountains. However, to the best of our current knowledge and after extensive monitoring, there is only a single locality for this species.

**Range description**

This is a single water tunnel endemic restricted to one spring water tunnel in the Judean mountains, Israel.

**Extent of occurrence (EOO) estimate (km^2^):** <1

**Trend:** Unknown.

**AOO (km^2^):** 4

**Trend:** Unknown.

**Number of locations: 1**

**Justification for number of locations:** only found in one spring water tunnel despite extensive surveys through the region.

**Habitat**

**System:** terrestrial

**Habitat specialist:** Yes

The species is a specialized troglobite living in the dark zone of ’En Sarig spring water tunnel, where temperature and humidity conditions are constant.

**Trend in extent, area or quality?:** Yes

**Justification for trend:**

Humidity and temperature change, changes in abiotic conditions due to extreme events such as global warming.

**Habitat importance:** Major Importance

**Habitats:**

- 5.9. Wetlands (inland) – Freshwater springs and oases

- 5.18. Wetlands (inland) – Karst and other subterranean hydrological systems (inland)

- 15.2 Ponds [below 8 ha]

- 15.10 Karst and Other Subterranean Hydrological Systems [human-made]

**Ecology**

**Size:** 6-8 mm

**Generation length (yr):** unknown

**Ecology and traits (narrative)**

The species builds funnel-webs in small holes in rock walls and base of the walls of the tunnel. Can be found mainly in the dark zones of the water tunnel (Aharon et al. 2023).

**Threats:**

This water tunnel is not protected by law and is very touristic during the summertime. Abiotic conditions within the water tunnel are affected by the visitors. Anthropogenic effects such as fire in the area near the water tunnel pose additional threats.

This species is dependent on the special conditions of humidity in the water tunnel. Extreme weather events such as drought and high temperatures could all destroy the water tunnel conditions in just one event and will have a severe effect on this micro-predator.

**Ongoing**

- 6. Human intrusions & disturbance

- 6.1. Human intrusions & disturbance - Recreational activities

- 7.1 Fire & Fire Suppression

- 7.1.1. Increase in Fire Frequency/Intensity: campfires and light inside the cave

- 11. Climate Change & Severe Weather

- 11.2 Extreme droughts

- 11.3 Temperature extremes

**Conservation**

As the water tunnel is not protected by law, conservation action is needed.

**Research needed:**

- 3. Monitoring

- 3.1. Monitoring - Population trends

- 3.4. Monitoring - Habitat trends

Critically Endangered (CR): B1a,b(iii,v)+B2a,b(iii,v)

3. *Tegenaria naasane* Aharon & Gavish-Regev, 2023

**Assessment Rationale**

This is a rare, endemic cave-dwelling funnel-web spider, found only in a limited area in the deep sections of A’rak Na’asane Cave (Z and K chambers in the cave map, see (Aharon et al., 2023; Frumkin et al., 2018)). Although these are two, separate chambers in one cave, they can be treated as one locality as all threats will affect both chambers. Extensive surveys in other chambers and other caves have been conducted and the species was not found elsewhere. The abiotic conditions in the cave, and especially in those two chambers with a steady temperature of 25.6–25.7 ◦C and a high relative humidity of 98.1–98.7 %, are fragile and may be threatened by climate-related changes and extreme weather events such as drought and heat, as well as anthropogenic effects such as fire in the cave. The micro-climate in the cave is very different from the desertic environment in which the cave is located, at the border between the semiarid eastern Samaria region arid northern Judean desert (Frumkin et al., 2018). In addition, this species and the whole food-web in the deep zones of the cave are dependent on guano produced by a colony of hundreds of insectivorous bat species *Asellia tridens* (´E. Geoffroy, 1813) seasonally inhabiting the cave, who themselves are threatened and in decline due to pesticide use in the past (Yom-Tov & Kadmon, 1998). The habitat extent and quality, and the number of adults are therefore projected to decline in the near future, making it Critically Endangered under Categories B1 and B2. Given the number of observations of very few individuals, less than 10 in total, despite extensive surveys, with very few adults it is very likely to be a very small population of less than 50 individuals in a single location, making this species Critically Endangered under Category D.

**Species information**

*Tegenaria naasane* Aharon & Gavish-Regev, 2023

**Common names**

Cave blind funnel-spider

**Taxonomy**

**Kingdom	Phylum	Class	Order	Family**

Animalia	Arthropoda	Arachnida	Araneae	Agelenidae

**Taxonomic notes**

This troglobitic species is found in deep sections of a single cave in the border between eastern Samaria and northern Judean desert (Palestinian authorities, West Bank) (Aharon et al., 2023). This spider species is eyeless, missing all ocelli and eye pigmentation (Figure 3), females and juveniles are highly depigmented.

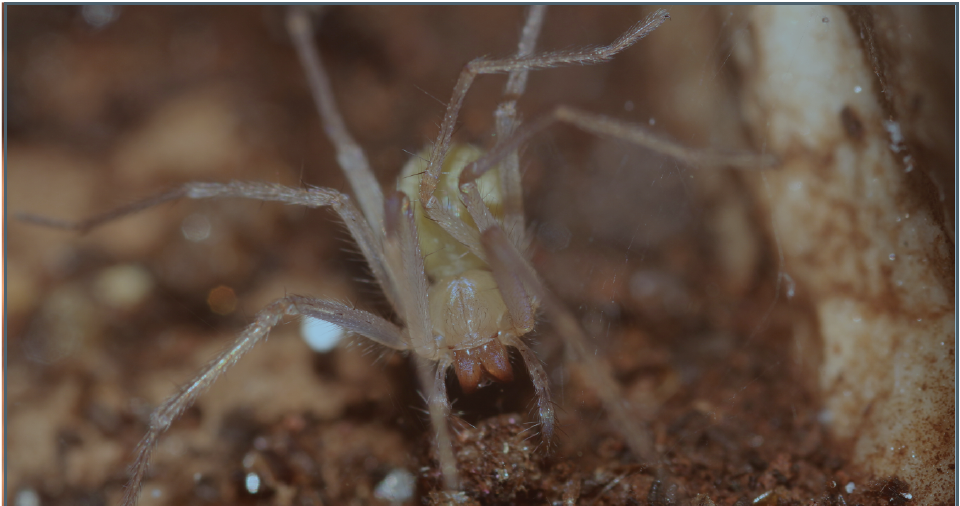

**Figure 3:** *Tegenaria naasane* Female,A’rak Na’asane Cave. Shlomi Aharon.

**Region for assessment:** Global

**Biogeographic realm:**  Palearctic, Levant

**Countries:** Palestine

**Map of records:**

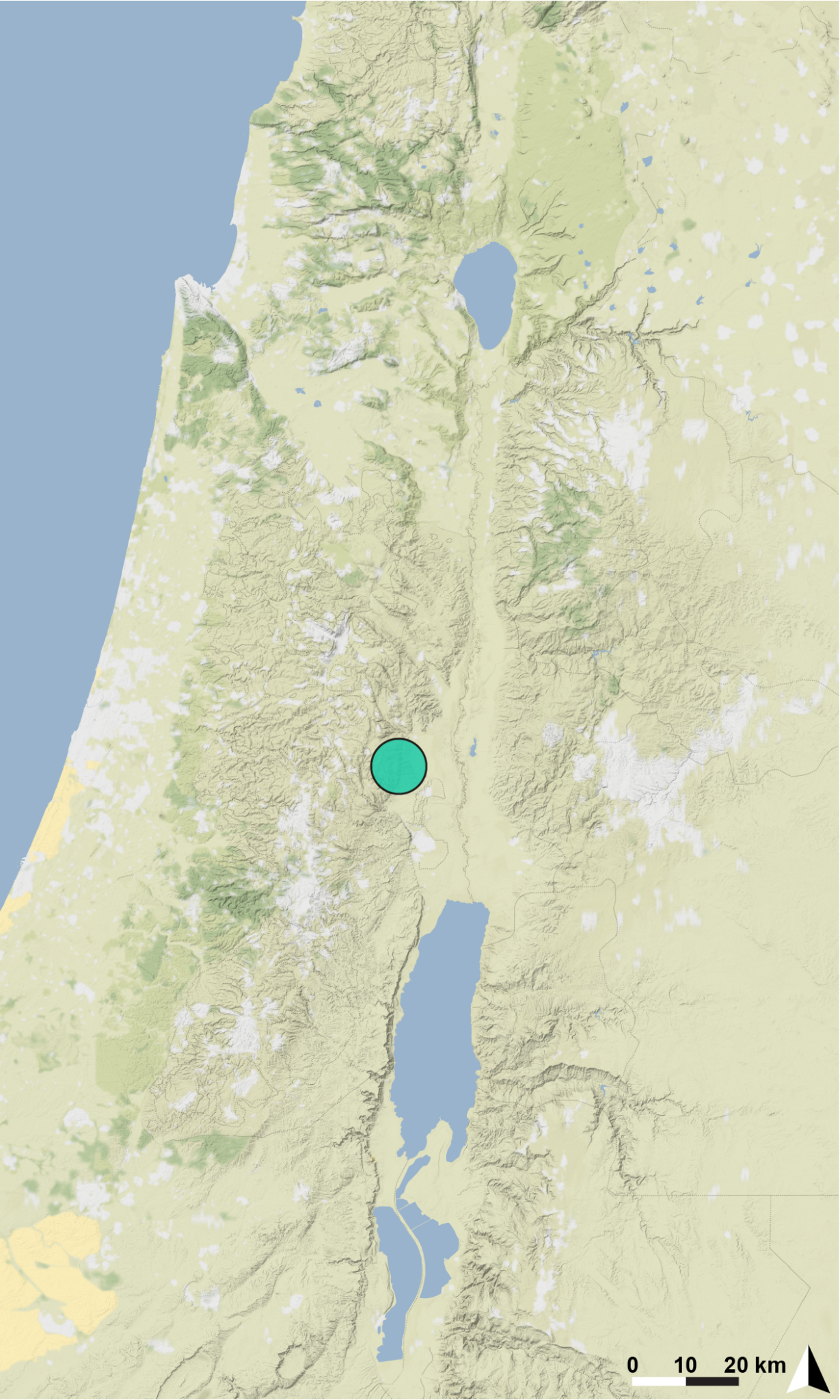

**Basis of EOO and AOO:** Observed

**Basis (narrative)**

Although more than 100 caves in Israel and Palestine were surveyed intensely for arachnids since 2012, A’rak Na’asane Cave is the only locality known for this anophthalmic *Tegenaria* species.

**Range description**

This is a single cave endemic restricted to a limited area in two deep chambers of A’rak Na’asane Cave (remote parts of Z and K chambers in the cave map, see (Aharon et al., 2023).

**Extent of occurrence (EOO) estimate (km^2^):** <1

**Trend:** Unknown.

**AOO (km^2^):** 4

**Trend:** Unknown.

**Number of locations: 1**

**Justification for number of locations:** only found in one cave despite extensive surveys through the region.

**Habitat**

**System:** terrestrial

**Habitat specialist:** Yes

The species is a specialized troglobite living in a limited area in the deep sections of a very large maze cave, A’rak Na’asane (length 2238 m) where temperature and humidity conditions are constant. It is restricted to two remote chambers with a steady temperature of 25.6–25.7 ◦C and a high relative humidity of 98.1–98.7 %.

**Trend in extent, area or quality?:** Yes

**Justification for trend:**

Humidity and temperature change, changes in abiotic conditions due to extreme events such as global warming. This species is dependent on bat guano in the cave. If the bat population declines this could wipe out the species. All bats are threatened in Israel due to pesticides and other threats. Fire risk is likely due to anthropogenic fires in the cave.

**Habitat importance:** Major Importance

**Habitats:**

- 7. Caves and Subterranean Habitats (non-aquatic)

- 7.1. Caves and Subterranean Habitats (non-aquatic) – Caves

**Ecology**

**Size:** 7.5 mm

**Generation length (yr):** unknown

**Ecology and traits (narrative)**

The species builds funnel-webs in small holes in walls of the cave, as well as under and near stones in the cave. It can be found at the edge of the guano deposition only in a limited area in two deep sections of the cave (Aharon et al. 2023).

**Threats:**

This cave is not protected and is used by locals for livestock, which significantly affects the conditions within the cave. In addition, climate change is predicted to affect the entire cave food-web and will have a severe effect on this micro-predator. Extreme weather events such as drought and high temperatures could all destroy the cave conditions in just one event. Anthropogenic effects such as fire and use of the cave for livestock resting is an additional threat. This species is dependent on the presence of guano from insectivorous bats in the cave, as it feeds on other arthropods that live and feed on the guano.

**Ongoing**

- 2. Agriculture

- 2.3 Livestock Ranching

- 2.3.1 Nomadic

- 2.3.2 Small-holder Ranching

- 6. Human intrusions & disturbance

- 6.1. Human intrusions & disturbance - Recreational activities

- 7.1 Fire & Fire Suppression

- 7.1.1. Increase in Fire Frequency/Intensity: campfires and light inside the cave

- 9. Pollution

- 9.3.3. Pesticides

- 11. Climate Change & Severe Weather

- 11.2 Extreme droughts

- 11.3 Temperature extremes

**Conservation**

The cave is not protected. In addition, bats are in decline which could severely reduce the resource availability for the species. The cave should be protected from anthropogenic fire and livestock rest. The deep chamber should be closed to visitors.

**Research needed:**

- 3. Monitoring

- 3.1. Monitoring - Population trends

- 3.4. Monitoring - Habitat trends

Critically Endangered (CR): B1a,b(iii)+B2a,b(iii), D

*4. Tegenaria ornit* Aharon & Gavish-Regev, 2023

**Assessment Rationale**

This is a rare, endemic cave-dwelling funnel-web spider, found only in one 95 m^2^ deep chamber of a single cave in the Karmel region. Extensive surveys in other chambers and other caves have been conducted and the species was not found elsewhere. The abiotic conditions in the cave are fragile and may be threatened by climate-related changes and extreme weather events such as drought and heat. In addition, conditions are threatened by anthropogenic effects including fire in the cave. This species and the whole food-web in the deep zone of the cave are dependent on guano produced by two insectivorous bat species (*Rhinolophus hipposideros minimus* Heuglin, 1861 and a *Myotis* sp.) seasonally inhabiting the cave, who themselves are threatened and in decline due to pesticides used in the past (Yom-Tov & Kadmon, 1998). The habitat extent and quality, and the number of adults are therefore projected to decline in the near future, making it Critically Endangered under Categories B1 and B2. Given the size of the chamber, and observations of less than 50 individuals, with very few adults and egg sacs (less than 10 of each were observed in two visits to the cave in 2020, and 2022) it is very likely to be a very small population of less than 50 individuals in a single location, making this species Critically Endangered under Category D.

**Species information**

*Tegenaria ornit* Aharon & Gavish-Regev, 2023

**Common names**

Cave blind funnel-spider

**Taxonomy**

**Kingdom	Phylum	Class	Order	Family**

Animalia	Arthropoda	Arachnida	Araneae	Agelenidae

**Taxonomic notes**

This troglobitic species is found in the deep chamber of a single cave in the Karmel mountain ridge in northern Israel (Aharon et al. 2023). Males, females and juveniles of this spider species are eyeless, missing all ocelli and eye pigmentation (Figure 4), females and juveniles are highly depigmented.

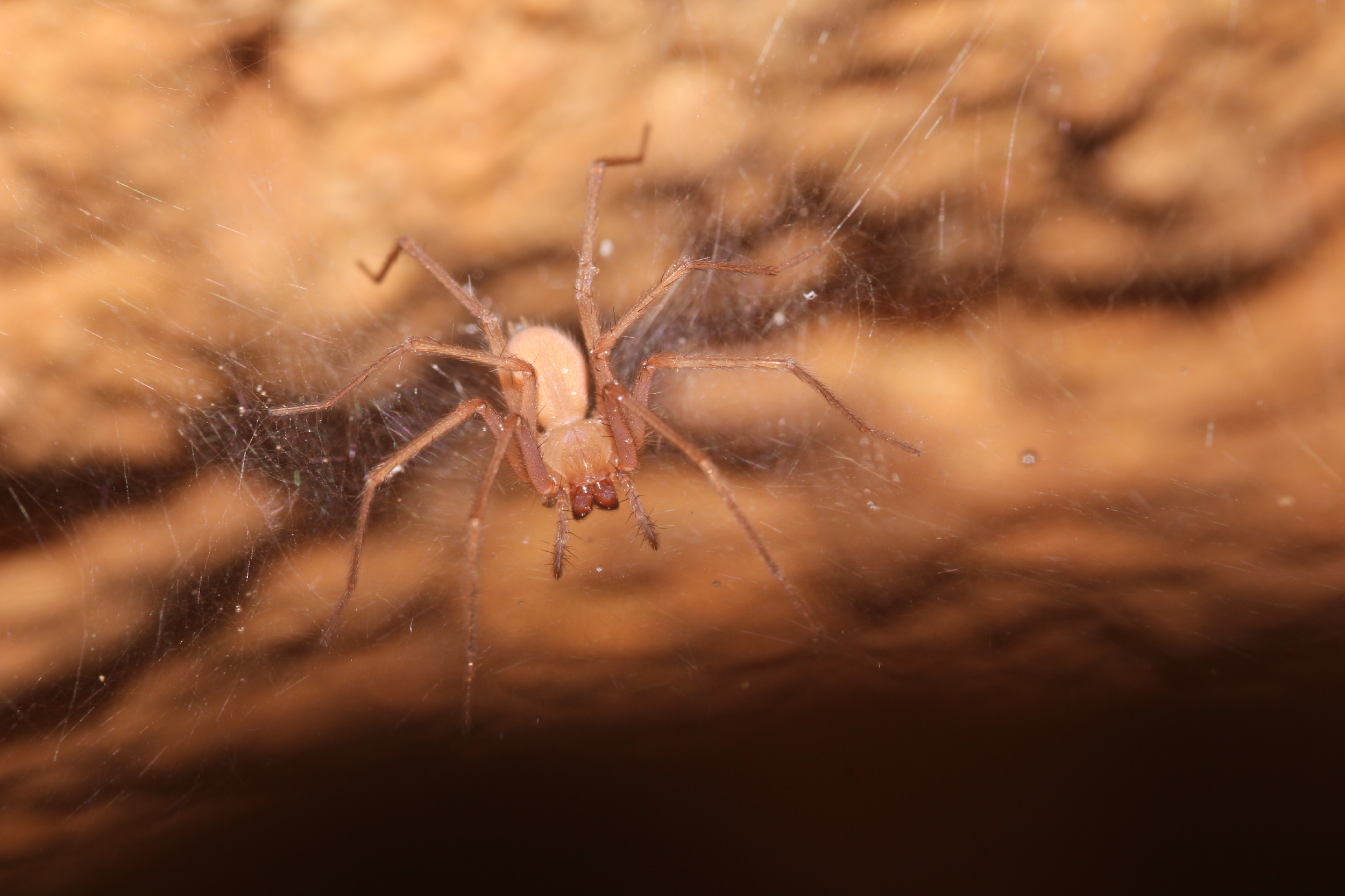

**Figure 4:** *Tegenaria ornit* Female, Ornit Cave. Shlomi Aharon.

**Region for assessment:** Global

**Biogeographic realm:**  Palearctic, Levant

**Countries:** Israel

**Map of records:**

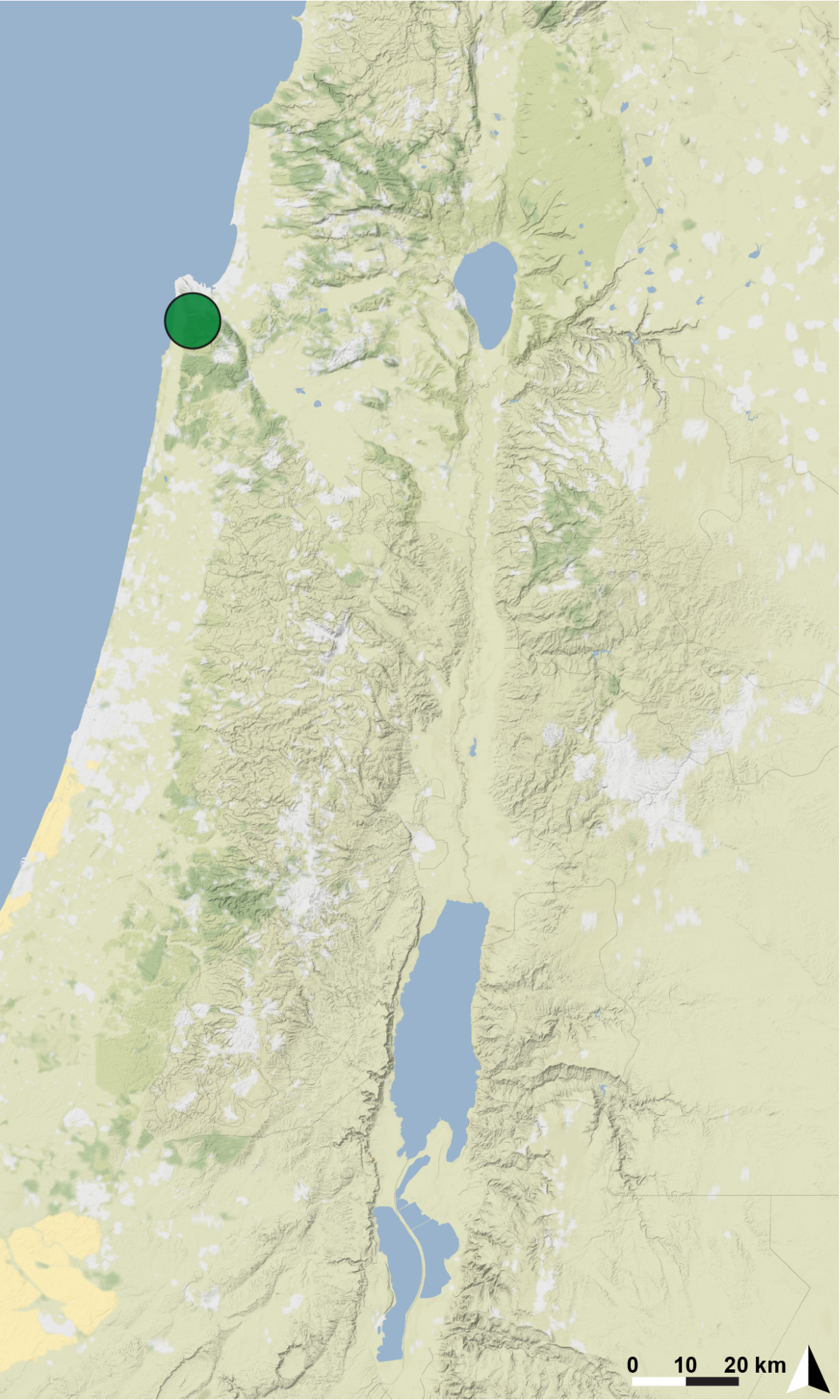

**Basis of EOO and AOO:** Observed

**Basis (narrative)**

Although more than 100 caves in Israel, including several caves in the Karmel mountains, were surveyed intensely for arachnids since 2012, Ornit Cave is the only locality known for this anophthalmic *Tegenaria* species.

**Range description**

This is a single-cave endemic restricted to the deep chamber (95 m^2^), of Ornit Cave, Karmel mountain ridge, Israel.

**Extent of occurrence (EOO) estimate (km^2^):** <1

**Trend:** Unknown.

**AOO (km^2^):** 4

**Trend:** Unknown.

**Number of locations: 1**

**Justification for number of locations:** only found in one cave despite extensive surveys through the region.

**Habitat**

**System:** terrestrial

**Habitat specialist:** Yes

The species is a specialized troglobite living in the deep chamber of one medium size cave, Ornit Cave, where temperature and humidity conditions are constant.

**Trend in extent, area or quality?:** Yes

**Justification for trend:**

Humidity and temperature change, changes in abiotic conditions due to extreme events such as global warming. This species is dependent on bat guano in the cave- if bat population declines this could wipe out the species. All bats are threatened in Israel due to pesticides and other threats. Fire risk is likely due to anthropogenic fire in the cave.

**Habitat importance:** Major Importance

**Habitats:**

- 7. Caves and Subterranean Habitats (non-aquatic)

- 7.1. Caves and Subterranean Habitats (non-aquatic) – Caves

**Ecology**

**Size:** 5-7 mm

**Generation length (yr):** unknown

**Ecology and traits (narrative)**

The species builds funnel-webs in small holes in rock walls of the cave, as well as under and near stones in the cave. Can be found only in the deep chamber (Aharon et al. 2023).

**Threats:**

This species is dependent on the presence of guano of insectivorous bats in the cave, as it feeds on other arthropods that live and feed on the guano. This cave is open to the public, and conditions within the cave are affected by the visitors. In addition, climate change is predicted to affect the abiotic conditions in the cave. Threats to the deep chamber food-web will have an especially severe effect on this micro-predator. Extreme weather events such as drought and high temperatures could destroy the cave conditions in just one event. Anthropogenic fire in the cave poses an additional threat.

**Ongoing**

- 6. Human intrusions & disturbance

- 6.1. Human intrusions & disturbance - Recreational activities

- 7.1 Fire & Fire Suppression

- 7.1.1. Increase in Fire Frequency/Intensity: campfires and light inside the cave

- 9. Pollution

- 9.3.3. Pesticides

- 11. Climate Change & Severe Weather

- 11.2 Extreme droughts

- 11.3 Temperature extremes

**Conservation**

The cave is currently protected by law, but bats are in decline which could severely reduce the resource availability for the species. The cave should be protected from anthropogenic fire. The deep chamber should be closed to visitors.

**Research needed:**

- 3. Monitoring

- 3.1. Monitoring - Population trends

- 3.4. Monitoring - Habitat trends

Critically Endangered (CR): B1a,b(iii,v)+B2a,b(iii,v), D

*5. Tegenaria trogalil* Aharon & Gavish-Regev, 2023

**Assessment Rationale**

This is an endemic cave-dwelling funnel-web spider, found in the twilight and deep zones of six caves in the upper Galilee mountains, northern Israel. Low variation in both morphology and DNA within cave populations, and higher variation between regions (eastern and western upper Galilee) and caves may suggest this is a species complex. Extensive surveys in other caves have been conducted and the species was not found elsewhere. The abiotic conditions in the caves (17.3–22.7 ◦C and 80–92.5 % humidity) are fragile and may be threatened by climate-related changes and extreme weather events such as drought and heat. In addition, conditions are threatened by anthropogenic effects including fire in the cave.This species and the whole food-web in the twilight and deep zones of some of those caves (Sharakh, Irav, Hutat Seter and Yir’on) are dependent on guano produced by the insectivorous bats (*Myotis* sp.), who themselves are threatened due to pesticide use in the past (Yom-Tov & Kadmon, 1998). Furthermore, some of those caves (Sharakh and Namer) are very touristic during the summertime. The habitat extent and quality, and the number of adults are therefore projected to decline in the near future, making it Vulnerable under Categories B1 and B2.

**Species information**

*Tegenaria trogalil* Aharon & Gavish-Regev, 2023

**Common names**

Cave funnel-spider

**Taxonomy**

**Kingdom	Phylum	Class	Order	Family**

Animalia	Arthropoda	Arachnida	Araneae	Agelenidae

**Taxonomic notes**

This troglobitic species is found in the twilight and deep zones of six caves in the upper Galilee, Israel (Aharon et al. 2023). Males, females and juveniles of this spider species are depigmented, and their eyes are reduced (Figure 5). Variation in morphological characters, especially eye reduction level and female epigynum, was observed between the caves, while high similarity was observed within specimens from the same cave. Six reduced eyes are usually present in Yir’on and Namer Cave populations, and eight reduced eyes are present at Sharakh cave population.

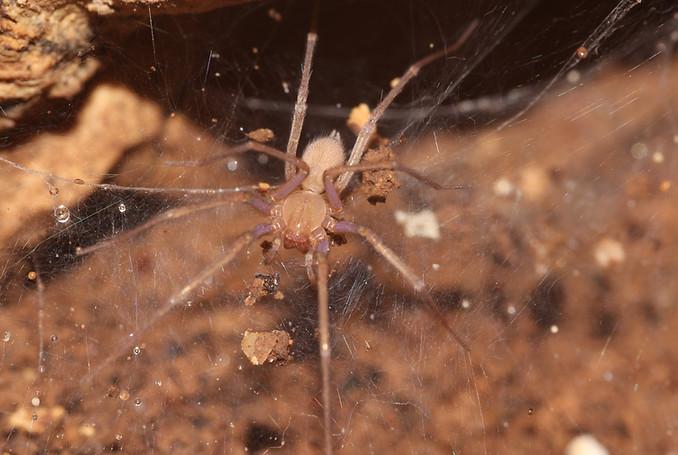

**Figure 5:** *Tegenaria trogalil* Female. Shlomi Aharon.

**Region for assessment:** Global

**Biogeographic realm:**  Palearctic, Levant

**Countries:** Israel

**Map of records:**

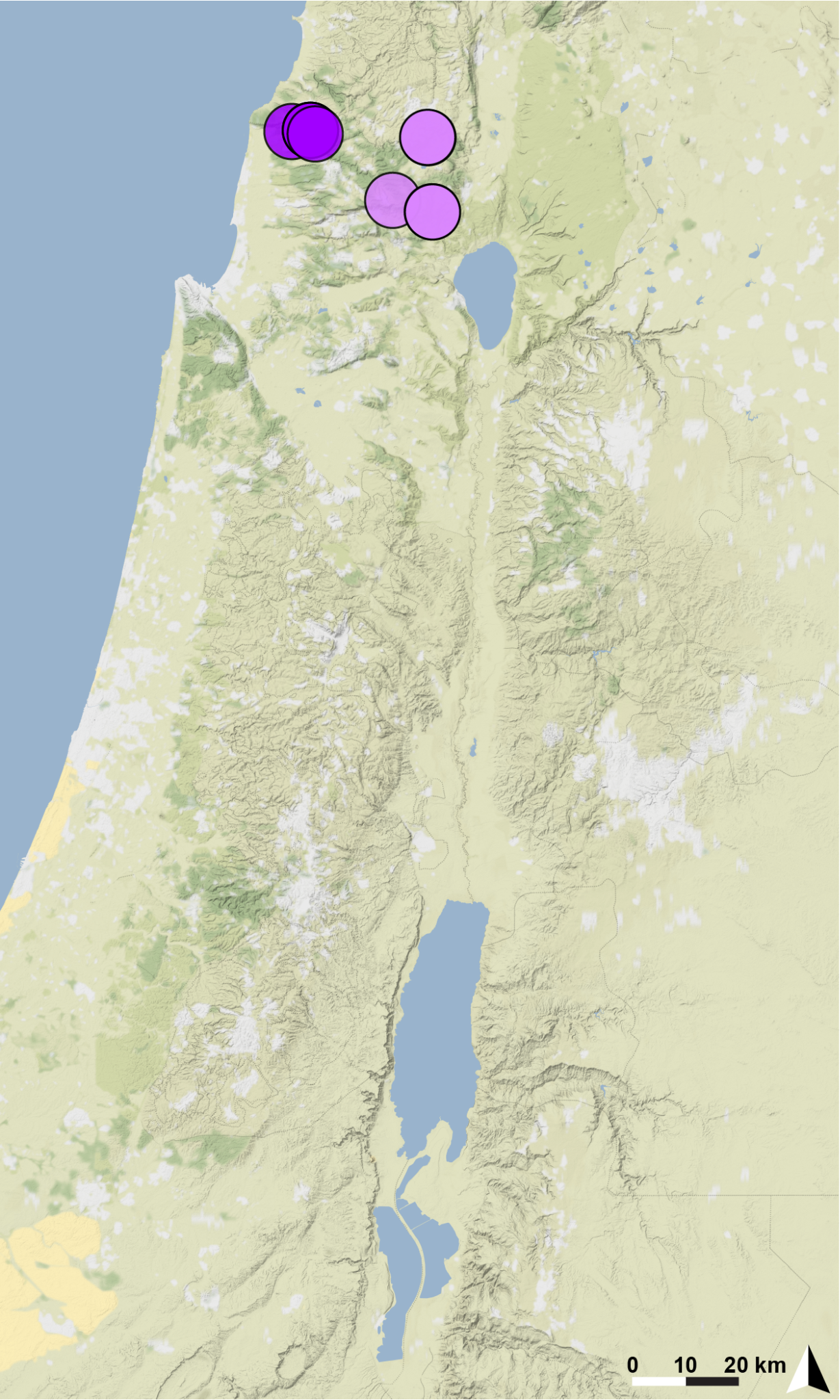

**Basis of EOO and AOO:** Observed

**Basis (narrative)**

Although more than 100 caves in Israel were surveyed intensely for arachnids since 2012, the upper Galilee caves Sharakh, Namer, Irav, Yir’on, Hutat Seter and Beit-Jann are the only localities known for this troglobitic *Tegenaria* species.

**Range description**

This is an endemic species restricted to the twilight and deep zones of six caves in The upper Galilee, Israel.

**Extent of occurrence (EOO) estimate (km^2^):** 212

**Trend:** Unknown.

**AOO (km^2^):** 24

**Trend:** Unknown.

**Number of locations:** 6

**Justification for number of locations:** only found in six caves despite extensive surveys through the region.

**Habitat**

**System:** terrestrial

**Habitat specialist:** Yes

The species is a specialized troglobite living in the twilight and deep zones of six caves in the upper Galilee, where temperature and humidity conditions are constant.

**Trend in extent, area or quality?:** Yes

**Justification for trend:**

Humidity and temperature change, changes in abiotic conditions due to extreme events such as global warming. This species is dependent on bat guano in some of the caves. If tbe bat population declines this could wipe out the species from those caves. All bats are threatened in Israel due to pesticides and other threats. Fire risk is likely due to manmade fire in some of the caves.

**Habitat importance:** Major Importance

**Habitats:**

- 7. Caves and Subterranean Habitats (non-aquatic)

- 7.1. Caves and Subterranean Habitats (non-aquatic) – Caves

**Ecology**

**Size:** 5-6.5 mm

**Generation length (yr):** unknown

**Ecology and traits (narrative)**

The species builds funnel-webs in small holes in rock walls of the caves, as well as under and near stones in the cave. Can be found only in the twilight and deep zones (Aharon et al. 2023).

**Threats:**

This species is found in the upper Galilee, and in some of these caves is dependent on the presence of guano of insectivorous bats, as it feeds on other arthropods that live and feed on the guano. These caves are open to the public, and conditions within the caves are affected by the visitors. In addition, climate change is predicted to affect the entire cave abiotic conditions and especially the deep zones food-web. Extreme weather events such as drought and high temperatures could all destroy the cave conditions in just one event and will have a severe effect on this micro-predator. In addition, anthropogenic effects such as fire in the caves pose an additional threat.

**Ongoing**

- 6. Human intrusions & disturbance

- 6.1. Human intrusions & disturbance - Recreational activities

- 7.1 Fire & Fire Suppression

- 7.1.1. Increase in Fire Frequency/Intensity: campfires and light inside the cave

- 9. Pollution

- 9.3.3. Pesticides

- 11. Climate Change & Severe Weather

- 11.2 Extreme droughts

- 11.3 Temperature extremes

**Conservation**

Only some of these caves are protected by law, but bats are in decline which could severely reduce the resource availability for the species. The caves should be protected from anthropogenic fire. The deep chambers should be close to visitors.

**Research needed:**

- 3. Monitoring

- 3.1. Monitoring - Population trends

- 3.4. Monitoring - Habitat trends

Vulnerable (VU): B1a,b(iii,v)+B2a,b(iii,v)

*6. Tegenaria yaaranford* Aharon Gavish-Regev, 2023

**Assessment Rationale**

This is an endemic cave-dwelling funnel-web spider, found only in the twilight and dark zones of a single cave in the western slope of the Judean mountains. Extensive surveys in other caves have been conducted and the species was not found elsewhere. The abiotic conditions in the cave are fragile and may be threatened by climate-related changes and extreme weather events such as drought and heat. In addition, conditions are threatened by anthropogenic effects including fire in the cave. In addition, the species and the whole food-web in the twilight and deep zones of the cave are dependent on guano produced by the frugivorous bat *Rousettus aegyptiacus* (Geoffroy, 1810), who themselves were threatened in Israel due to pesticide use (Korine et al., 1999). Furthermore, this cave is highly touristic and can receive more than 600 visitors a day during the summertime. The habitat extent and quality, and the number of adults are therefore projected to decline in the near future, making it Critically Endangered under Categories B1 and B2.

**Species information**

*Tegenaria yaaranford* Aharon Gavish-Regev, 2023

**Common names**

Cave funnel-spider

**Taxonomy**

**Kingdom	Phylum	Class	Order	Family**

Animalia	Arthropoda	Arachnida	Araneae	Agelenidae

**Taxonomic notes**

This troglobitic species is found in the twilight and dark zones of a single cave in the western slope of the Judean mountains in Israel (Aharon et al. 2023). Males, females and juveniles of this spider species are highly depigmented and bare eight highly reduced eyes, (Figure 6).

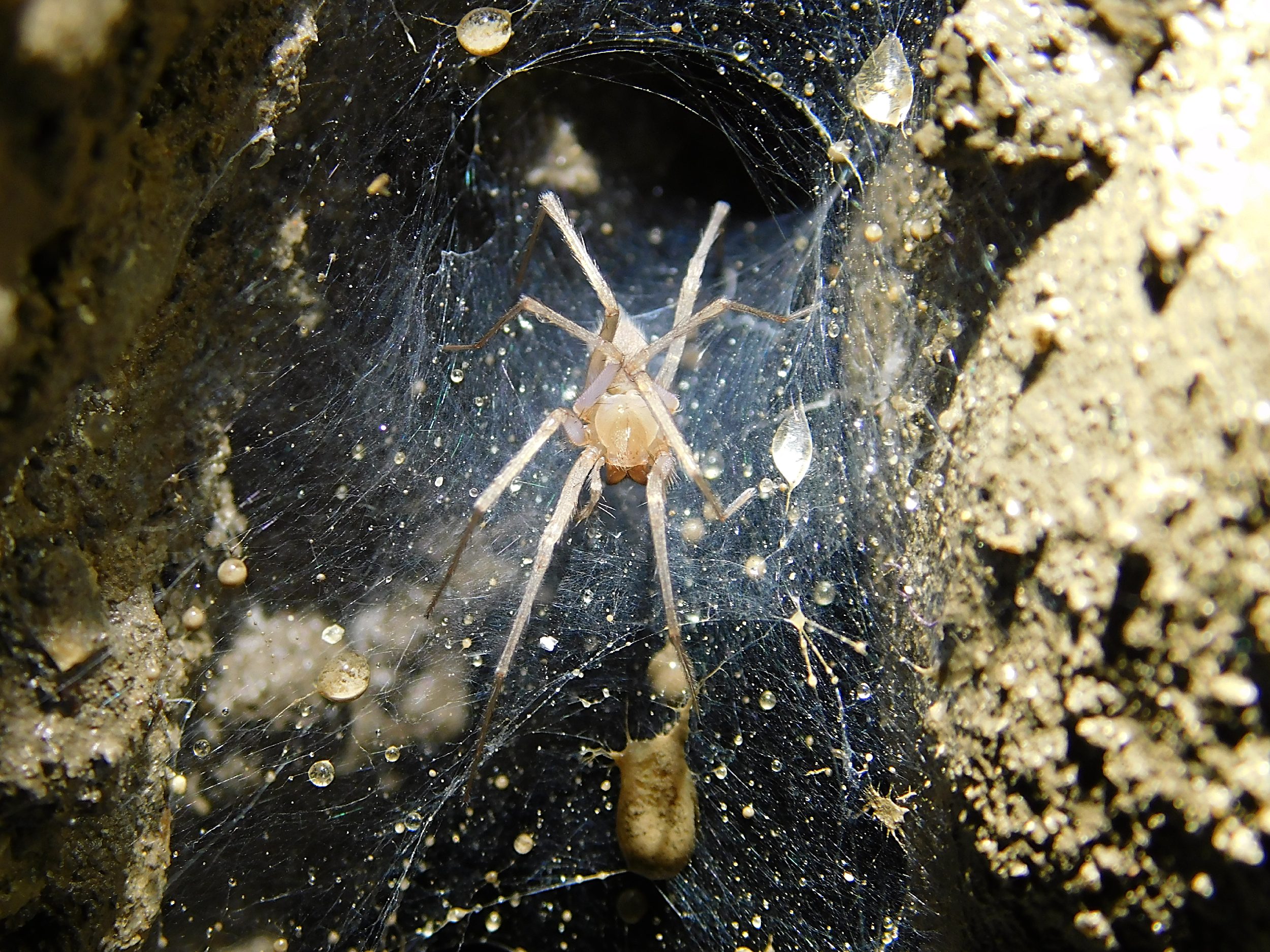

**Figure 6:** *Tegenaria yaaranford* Female, Te’omim Cave. Shlomi Aharon.

**Region for assessment:** Global

**Biogeographic realm:**  Palearctic, Levant

**Countries:** Israel

**Map of records:**

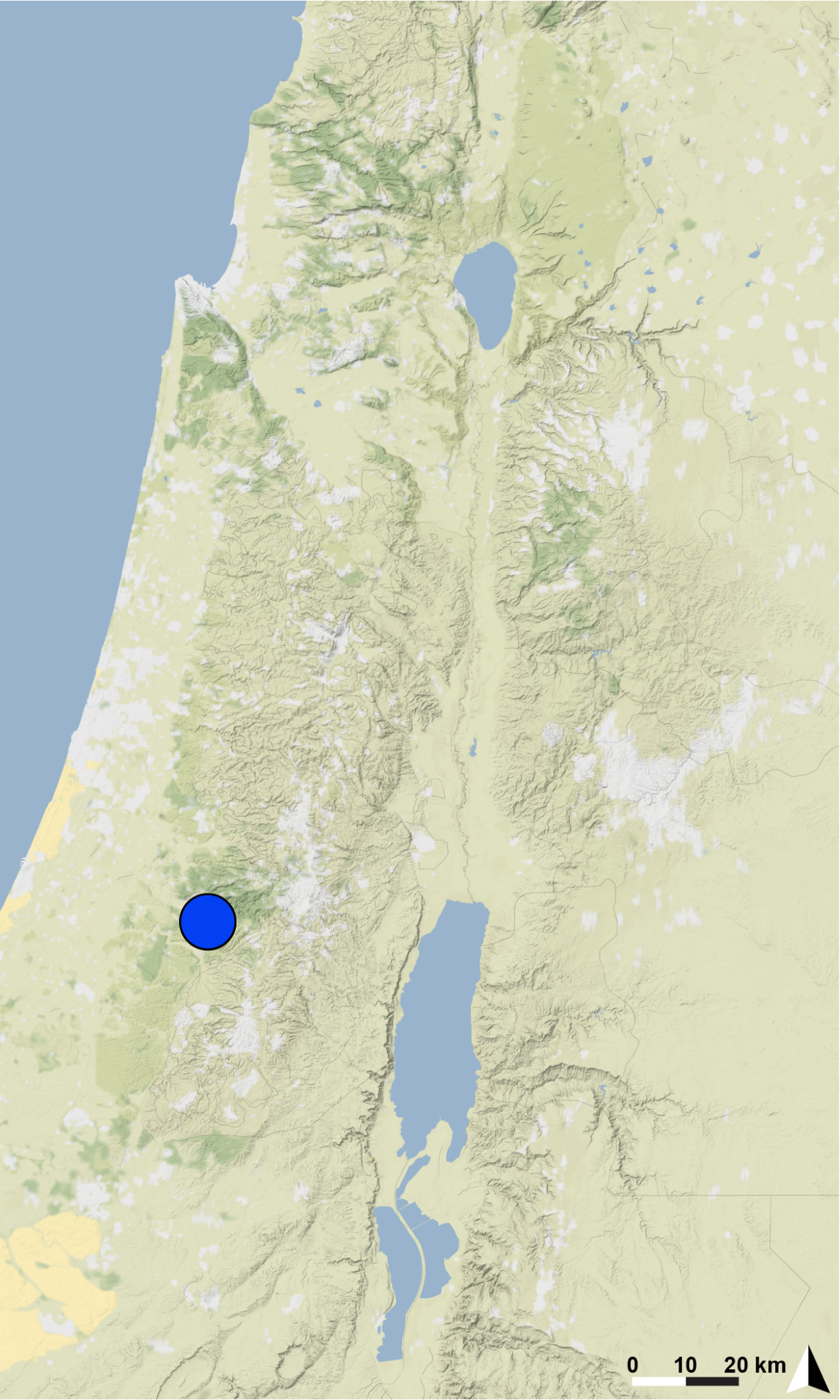

**Basis of EOO and AOO:** Observed

**Basis (narrative)**

Although more than 100 caves in Israel, including several caves in the Judean mountains, were surveyed intensely for arachnids since 2012, Te’omim Cave is the only locality known for this troglobitic *Tegenaria* species.

**Range description**

This is a single cave endemic restricted to the twilight and dark zones of Te’omim Cave, the Judean mountains, Israel. The large hall of the twilight zone of the cave is 50X70 m, (about 3500m2 volume)

**Extent of occurrence (EOO) estimate (km^2^):** <1

**Trend:** Unknown.

**AOO (km^2^):** 4

**Trend:** Unknown.

**Number of locations: 1**

**Justification for number of locations:** only found in one cave despite extensive surveys through the region.

**Habitat**

**System:** terrestrial

**Habitat specialist:** Yes

The species is a specialized troglobite living in the twilight and dark zones of Te’omim Cave, where temperature and humidity conditions are constant.

**Trend in extent, area or quality?:** Yes

**Justification for trend:**

Humidity and temperature change, changes in abiotic conditions due to extreme events such as global warming. This species is dependent on bat guano in the cave. If the bat population declines this could wipe out the species. All bats are threatened in Israel due to pesticides and other threats. Fire risk is likely due to anthropogenic fire in the cave. Today, Te’omim Cave is open to tourists from March to November although the nature reserve authorities close the cave between November and March.

**Habitat importance:** Major Importance

**Habitats:**

- 7. Caves and Subterranean Habitats (non-aquatic)

- 7.1. Caves and Subterranean Habitats (non-aquatic) – Caves

**Ecology**

**Size:** 4-9 mm

**Generation length (yr):** unknown

**Ecology and traits (narrative)**

The species builds funnel-webs in small holes in rock walls of the cave, as well as under or near stones in the cave. Can be found only in the twilight and dark zones of the cave (Aharon et al. 2023).

**Threats:**

This species is dependent on the presence of guano of frugivorous bats in the cave, as it feeds on other arthropods that live and feed on the guano. This cave is a touristic cave open to the public from March to November. Tourism may affect the abiotic conditions in the cave due to changes in temperature, humidity and oxygen saturation. In addition, climate change is predicted to affect the entire cave abiotic conditions and will have a severe effect on this micro-predator. Extreme weather events such as drought and high temperatures could all destroy the cave conditions in just one event. Anthropogenic fire in the cave is an additional threat.

**Ongoing**

- 6. Human intrusions & disturbance

- 6.1. Human intrusions & disturbance - Recreational activities

- 7.1 Fire & Fire Suppression

- 7.1.1. Increase in Fire Frequency/Intensity: campfires and light inside the cave

- 9. Pollution

- 9.3.3. Pesticides

- 11. Climate Change & Severe Weather

- 11.2 Extreme droughts

- 11.3 Temperature extremes

**Conservation**

The cave is currently protected by law, but it is highly touristic and can receive more than 600 visitors a day during the summertime, and therefore light is another concern, although it is closed for visitors during the winter by the nature reserve authorities.

**Research needed:**

- 3. Monitoring

- 3.1. Monitoring - Population trends

- 3.4. Monitoring - Habitat trends

Critically Endangered (CR): B1a,b(iii,v)+B2a,b(iii,v)

*7. Tegenaria yotami* Gavish-Regev & Aharon, 2023

**Assessment Rationale**

This is a rare, endemic cave-dwelling funnel-web spider, found only in the isolated dark parts of a single touristic cave in the western slope of the Judean mountains. Extensive surveys in other caves have been conducted and the species was not found elsewhere. The abiotic conditions in the cave are fragile and may be threatened by climate-related changes and extreme weather events such as drought and heat. In addition, the touristic nature of the cave may affect its abiotic conditions. The habitat extent and quality, and the number of adults are therefore projected to decline in the near future, making it Critically Endangered under Categories B1 and B2. Given the observations of very few individuals (less than 10, no males were observed) it is very likely to be a very small population of less than 50 individuals in a single location, making this species Critically Endangered under Category D.

**Species information**

*Tegenaria yotami* Gavish-Regev & Aharon, 2023

**Common names**

Cave funnel-spider

**Taxonomy**

**Kingdom	Phylum	Class	Order	Family**

Animalia	Arthropoda	Arachnida	Araneae	Agelenidae

**Taxonomic notes**

This troglobitic species is found in the dark and isolated parts of a single cave in the western slope of the Judean mountains in Israel (Aharon et al. 2023). Females and juveniles of this spider species are highly depigmented and bare six highly reduced eyes, lacking the anterior median eyes (Figure 7), males are unknown.

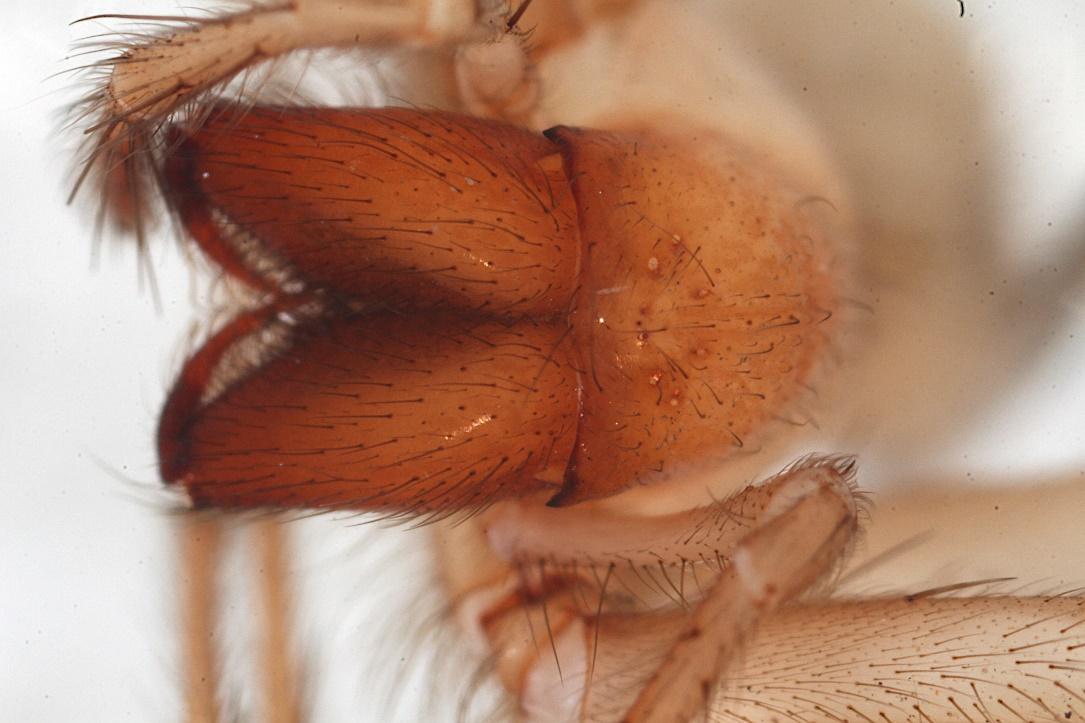

**Figure 7:** *Tegenaria yotami* Female, Soreq Cave. Shlomi Aharon.

**Region for assessment:** Global

**Biogeographic realm:**  Palearctic, Levant

**Countries:** Israel

**Map of records:**

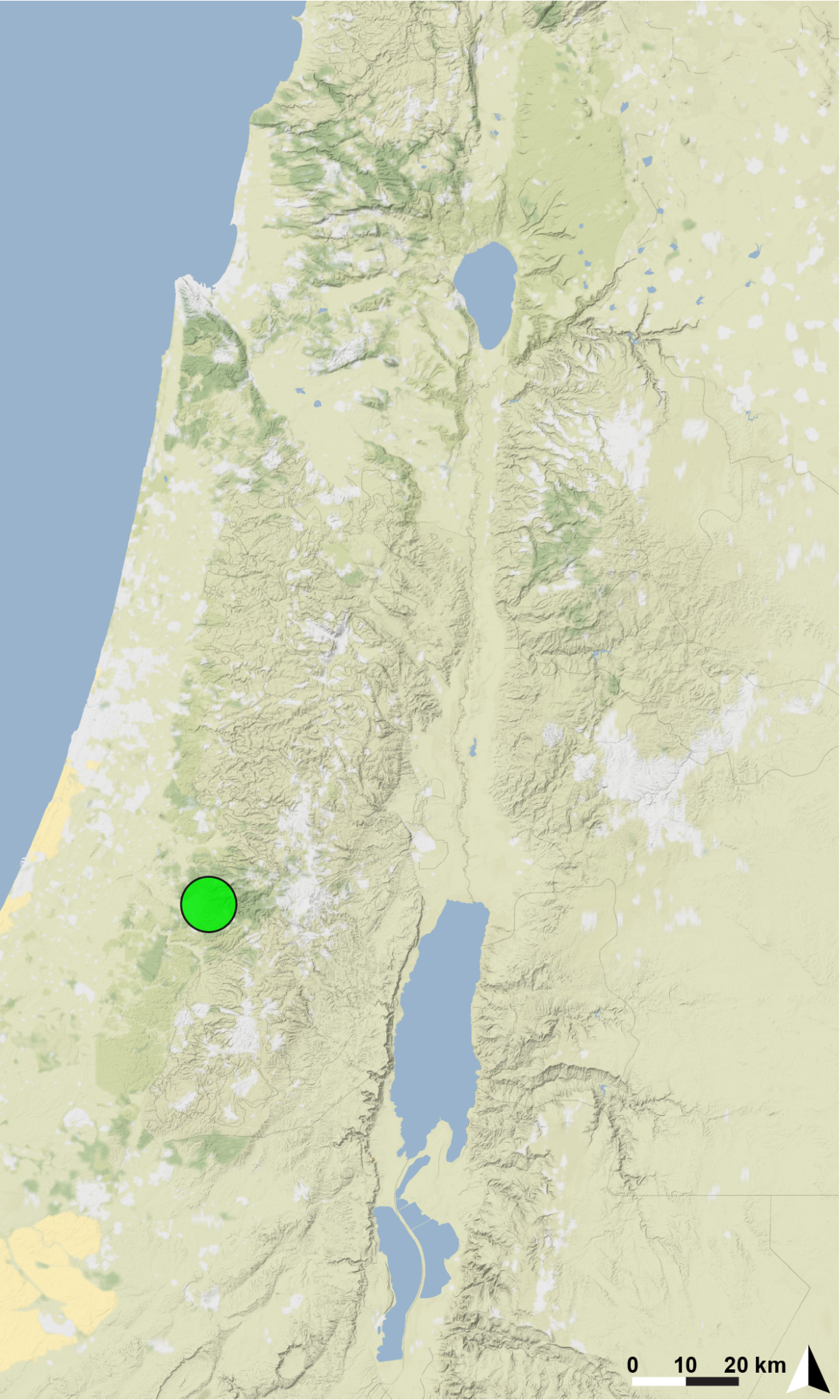

**Basis of EOO and AOO:** Observed

**Basis (narrative)**

Although more than 100 caves in Israel, including several caves in the Judean mountains, were surveyed intensely for arachnids since 2012, Soreq Cave is the only locality known for this troglobitic *Tegenaria* species.

**Range description**

This is a single-cave endemic restricted to the dark and isolated parts of Soreq Cave, the Judean mountains, Israel. Although the cave is large (4800 m2 volume), the spiders were found in restricted areas in the cave.

**Extent of occurrence (EOO) estimate (km^2^):** <1

**Trend:** Unknown.

**AOO (km^2^):** 4

**Trend:** Unknown.

**Number of locations: 1**

**Justification for number of locations:** only found in one cave despite extensive surveys through the region.

**Habitat**

**System:** terrestrial

**Habitat specialist:** Yes

The species is a specialized troglobite living in the dark and isolated parts of Soreq touristic cave, where temperature and humidity conditions are constant.

**Trend in extent, area or quality?:** Yes

**Justification for trend:**

Humidity and temperature change, changes in abiotic conditions due to extreme events such as global warming. Today, Soreq Cave is open to tourists on a daily basis, although the nature reserve authorities are making an effort to close the cave tightly between visits (from the afternoon to the next morning).

**Habitat importance:** Major Importance

**Habitats:**

- 7. Caves and Subterranean Habitats (non-aquatic)

- 7.1. Caves and Subterranean Habitats (non-aquatic) – Caves

**Ecology**

**Size:** 6-7 mm

**Generation length (yr):** unknown

**Ecology and traits (narrative)**

The species builds funnel-webs in small holes in rock walls of the cave, as well as under or near stones in the cave. Can be found only in the dark and isolated parts of the cave (Aharon et al. 2023).

**Threats:**

This cave is open to the public, and conditions within the cave are affected by the visitors. Tourism may have an impact on cave conditions, such as changes in temperature, humidity and oxygen saturation, as well as on the species assemblage in the cave. We recently found in the cave an invasive spider species (*Eidmannella pallida* Emerton, 1875). This invasive spider species was found in another cave (Ayyalon Cave), while a local endemic pseudoscorpion disappeared in parallel with the appearance of the invasive species (Aharon, unpublished observation). In addition, climate change is predicted to affect the abiotic conditions of the cave and will have a severe effect on this micro-predator. Extreme weather events such as drought and high temperatures could all destroy the cave conditions in just one event.

**Ongoing**

- 6. Human intrusions & disturbance

- 6.1. Human intrusions & disturbance - Recreational activities

- 8. Invasive & Other Problematic Species, Genes & Diseases

- 8.1. Invasive Non-Native/Alien Species/Diseases

- 8.1.2. *Eidmannella pallida* Emerton, 1875

- 11. Climate Change & Severe Weather

- 11.2 Extreme droughts

- 11.3 Temperature extremes

**Conservation**

The cave is currently protected by law, but although the nature reserve authorities are making an effort to close the cave tightly between visits, it has a high volume of visitors. Therefore, light is being used during opening hours, and the cave is open during activity times. Rodents invading the cave were also observed.

**Research needed:**

- 3. Monitoring

- 3.1. Monitoring - Population trends

- 3.4. Monitoring - Habitat trends

Critically Endangered (CR): B1a,b(iii,v)+B2a,b(iii,v), D
