## Supplemental File 2 for "Machine learning approaches to assess microendemicity and conservation risk in cave-dwelling arachnofauna"

S*pecies sampling and sequencing methods*

We sequenced samples from 161 *Tegenaria* specimens freshly collected from field campaigns in Israel (2018-2020). The previously extracted samples were eluted in EDTA, which necessitated an initial bead cleanup due to its interaction with reagents downstream in the enzymatic fragmentation step of library preparation. The remaining samples were extracted from specimens using a Qiagen DNeasy Blood & Tissue Kit and eluted in 10 mM Tris-HCl. We added AMPure XP beads (Beckman Coulter) directly to 1.5 mL microcentrifuge tubes containing the pre-extracted samples and incubated the mixture for 10 minutes at room temperature. We then removed the supernatant and performed 2 washes of the beads with 80% EtOH. We subsequently resuspended the beads in 52 µL of 10 mM Tris-HCl, incubated the mixture for 2 minutes, and transferred the sample to a fresh 1.5 mL microcentrifuge tube. Samples were then quantified using a Qubit High Sensitivity dsDNA Assay Kit and Qubit Fluorometer. Next, we used a KAPA HyperPlus Kit to perform enzymatic fragmentation, end repair/A-tailing, adapter ligation, and library amplification following the KAPA Library Construction Protocol v5.19. For adapters, we used universal iTru stubs (EHS DNA Laboratory, University of Georgia). 5 µL of iTru stubs at 5µM concentration were added to the end repair and A-tailing reaction product (30 µL) along with 15 µL of ligation buffer and 5 µL of DNA ligase. We repeated the bead cleanup procedure outlined before, this time using KAPA HyperPure beads. We then quantified the libraries again using a Qubit Fluorometer. Subsequently, we performed library amplification using dual index primers (i5 and i7) for eventual multiplex sequencing. Libraries were quantified once more after this step. We pooled samples by groups of eight, each with 125 ng of DNA for subsequent target capture. We followed the myBaits protocol v5.02 (Arbor Biosciences) for targeted enrichment using a spider-specific probe set (Kulkarni et al. 2020). Pools were then sent to the University of Wisconsin Biotechnology Center for sequencing. Sequencing was performed on an Illumina NovaSeq 6000 2 ´ 150 bp S1 flow cell.

To clean and trim our raw reads we used illumiprocessor v2.0. We employed the software package PHYLUCE v1.6 for subsequent data processing and analysis (Faircloth 2015). For quality control, we ran the script “phyluce_assembly_get_fastq_lengths” to generate summary statistics on our reads. To assemble the reads we ran “phyluce_assembly_assemblo_abyss” with ABySS v2.1.4. We performed another quality control check by running “phyluce_assembly_get_fasta_lengths” which generated summary statistics on the assembled contigs. Next came finding the actual UCE loci which utilized the script “phyluce_assembly_match_contigs_to_probes” to identify contigs that represented UCE loci in our probe set. To see how many UCE loci were recovered for each library (taxon), we made a configuration file containing our entire dataset and then ran “phyluce_assembly_get_match_counts.” This generated a data matrix configuration file from which FASTA data could be extracted for each taxon. Running the script “phyluce_assembly_get_fastas_from_match_counts,” we extracted the FASTA data from the loci for each taxon in the configuration file. We ran “phyluce_align_seqcap_algin” with MAFF[HS8] T v7.490 to align the loci. Then we ran edge trimming on the alignments with “phyluce_align_get_gblocks_trimmed_alignments_from_untrimmed” using Gblocks v0.91b[HS9] . We removed the locus names from the alignments by running “phyluce_align_remove_locus_name_from_files” which left us with alignments labeled only by taxon. To get summary statistics on the alignments and data matrices we ran “phyluce_align_get_align_summary_data.” To analyze a 50% complete data matrix, we ran “phyluce_align_get_only_loci_with_min_taxa” and selected a value of 0.50 for the “percentage complete” parameter. This narrowed our data to alignments that only appeared in 50% of taxa and moved these alignments to a new directory. To prepare for downstream phylogenomic analysis we ran “phyluce_align_concatenate_alignments” to generate a phylip file.
